# Testing Darwin’s naturalization conundrum based on taxonomic, phylogenetic and functional dimensions of vascular plants

**DOI:** 10.1101/847442

**Authors:** Jesús N. Pinto-Ledezma, Fabricio Villalobos, Peter B. Reich, Daniel J. Larkin, Jeannine Cavender-Bares

## Abstract

Charles Darwin posited two alternative hypotheses to explain the success of nonnative species based on their relatedness to incumbent natives: coexistence between them should be (i) more likely with greater relatedness (due to trait similarity that correlates with better matching to the environment), or (ii) less likely (due to biotic interference, such as competition). The paradox raised by the opposing predictions of these two hypotheses has been termed ‘Darwin’s naturalization conundrum’ (DNC). Using plant communities measured repeatedly over a 31-year time span across an experimental fire gradient in an oak savanna (Minnesota, USA) we evaluated the DNC by explicitly incorporating taxonomic, functional and phylogenetic information. Our approach was based on ‘focal-species’ such that the taxonomic, functional and phylogenetic structure of species co-occurring with a given nonnative species in local communities was quantified. We found three main results: first, nonnatives colonizers tended to co-occur most with closely related incumbent natives in recipient communities, except in the extreme ends of the fire gradient (i.e., communities with no fire and those subjected to high fire frequencies); second, with increasing fire frequency, nonnative species were functionally more similar to native species in recipient communities; third, functional similarity of co-occurring nonnatives and natives in recipient communities showed a consistent pattern over time, but the phylogenetic similarity shifted over time, suggesting that external forces (e.g., climate variability) are also relevant in driving the phylogenetic relatedness of nonnatives to natives in invaded communities. Our results provide insights for understanding the invasion dynamics across environmental gradients and highlight the importance of evaluating different dimensions of biodiversity in order to produce more powerful evaluations of species co-occurrence at different spatial and temporal scales.

## Introduction

The assembly and maintenance of ecological communities is a dynamic process operating over multiple spatial and temporal scales, that incorporates local niche-based interactions and sorting to stochastic and historical processes that may operate over large spatial scales (Tilman 2004, Cavender-Bares et al. 2009, 2018a, Pinto-Ledezma et al. 2019). Over the past millennium, human activities have greatly influenced these natural processes, through habitat degradation and biological invasions by moving species out of their native ranges, with negative consequences for biodiversity, ecosystem functioning, and human well-being (Sax et al., 2007, Thuiller et al. 2010, Vilà et al. 2011, Simberloff et al. 2013, Capinha et al. 2015).

Given the importance of biological invasions in determining current community structure (Pearson et al. 2018), understanding the causes of invasion success have become a major goal in ecology, evolution and conservation (Dawson et al. 2017). While there are many competing hypotheses for the success and failure of colonizing species (Blumenthal 2005, Jeschke et al. 2012, Jeschke 2014, Prins and Gordon 2014), two major hypotheses have been proposed as explanations for species invasion success that incorporate evolutionary relatedness as a primary consideration (Gallien and Carboni 2017, Ma et al. 2016, Cadotte et al. 2018). First, Darwin’s naturalization hypothesis (DNH; Box 1: Fig. 1A) suggests that nonnative species closely related to resident natives are less likely to invade native assemblages because the niches they could invade are already occupied by ecologically similar relatives (Daehler 2001). In contrast, the pre-adaptation hypothesis (PAH; Box 1: Fig. 1B) postulates that nonnative species closely related to resident natives should be favored precisely because of their niche similarity with native species, sharing traits that make them well-suited to the novel range (Ricciardi and Mottiar, 2006). Accordingly, the extent to which nonnative species are closely or distantly related to resident species may teach us whether competitive interactions or environmental filters, respectively, are dominant factors determining invasion success (Gallien and Carboni 2017, Cadotte et al. 2018). These opposing hypotheses both trace back to Darwin (1859) and together comprise ‘Darwin’s naturalization conundrum’ (DNC, Diez et al. 2008, Thuiller et al. 2010, Cadotte et al. 2018).

**Fig. 1.**
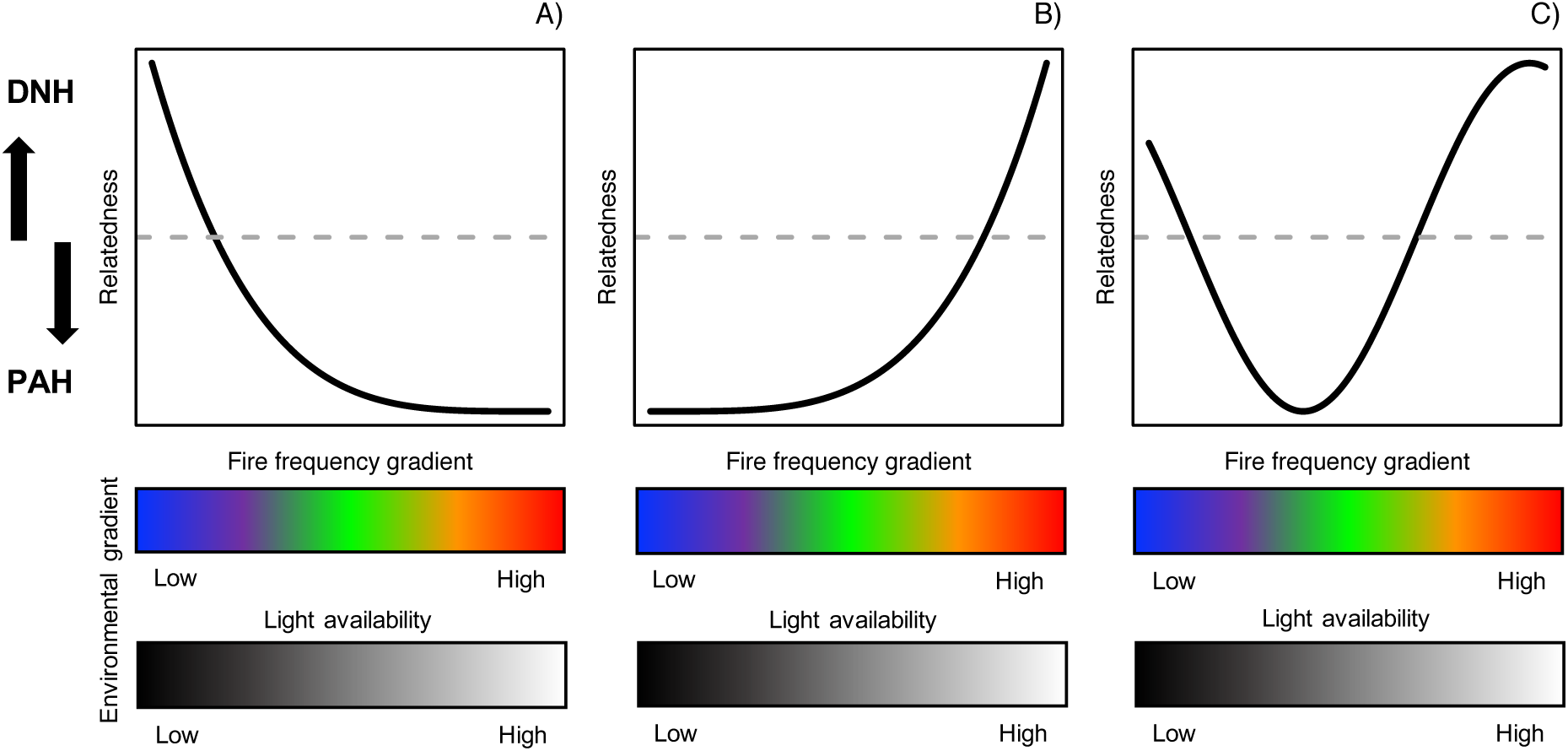
Hypotheses and predictions for co-occurrence patterns of focal-nonnative species with plants in their surrounding communities across environmental gradients at Cedar Creek. The curves depict theoretical expectations of changes in phylogenetic/functional structure across fire and light-availability gradients, explained in the main text.

Deciphering the connection between ecological and evolutionary processes in driving species distributions and the assembly of communities is crucial to understand the invasion success of nonnative species in recipient communities (Gallien and Carboni 2017, Cadotte et al. 2018, Pearson et al. 2018). Although the DNC represents an integrative explanation that links both ecological and evolutionary processes (reviewed in Cadotte et al. 2018), invasion is a dynamic process, i.e., nonnative species are continually expanding or retracting their geographical ranges across the regions they have recently colonized (Sax et al. 2007, Blackburn et al. 2015, Pannell 2015). Thus, the presence of nonnative species in a community does not necessarily indicate that they are optimally-adapted to the new environmental or niche conditions. One potential explanation for the spread of invasive species is the ‘ecological fitting hypothesis’ (EFH, Janzen 1985), which suggests that widespread species can occupy new places or environmental conditions without being perfectly adapted to them (Janzen 1985, Cavender-Bares et al. 2018b, but see Odour et al. 2016). In addition, functional traits underlie composition, community assembly and ecosystem processes (Cavender-Bares et al. 2009, Cavender-Bares et al. 2016, Lavorel et al. 2011, Reich 2014, Catford et al. 2019); thus, different functional traits or trait combinations can modulate the degree to which nonnative species are able to colonize and further adapt to the ecological conditions found in recipient communities (Blumenthal 2005, van Kleunen et al. 2010, Carboni et al. 2018; Catford et al 2019). Recent evidence suggests that successful invasive species tend to have higher values for traits associated with resource acquisition, dispersal, and establishment and competitive ability than local native species (van Kleunen et al. 2010, Carboni et al. 2018, Catford et al. 2019), indicating that they have similar or higher performance in the novel range than native species (Sax et al. 2007, Odour et al. 2016, but see González-Muñoz et al. 2014).

Several studies have evaluated the DNC across different spatial scales and systems (for a review see Cadotte et al. 2018, Gallien and Carboni 2017, Ma et al. 2016). However, few studies have explored the dynamics of species composition and relatedness within communities during the invasion process (Blackburn et al. 2015, Li et al. 2015) or the role of functional traits in modulating colonization and establishment by nonnative species (Marx et al. 2015, Carboni et al. 2018; but see Catford et al 2019). Although these studies have generally found similar results— from a phylogenetic perspective, nonnative species tend to coexist more with their close relatives (e.g., Li et al. 2015, Marx et al. 2015, Kusumoto et al. 2019)—the incorporation of functional information into analyses provides new insights regarding functional differentiation between coexisting species (Cavender-Bares et al. 2009, Cadotte et al. 2018); and consequently a way forward to understand how species’ ecological differences regulate the colonization, establishment and persistence of nonnative species within local native communities across spatial and temporal scales.

Here, using plant communities sampled over decades across an experimental fire gradient at Cedar Creek Ecosystem Science Reserve (hereafter Cedar Creek) in Minnesota, USA, we evaluate the DNC while explicitly incorporating taxonomic, functional, and phylogenetic information into our analyses. To do so, we apply a novel approach based upon the framework of Villalobos et al. (2013, 2017), extending the concept of species’ *functional/phylogenetic fields*— the overall functional/phylogenetic structure within a given species’ geographical range—to describe the functional and phylogenetic structure of species co-occurring with a focal species; for simplicity, we call this approach ‘focal-species’ (Box 1). In this approach, each species within a community is in turn selected as a focal species; and its phylogenetic/functional distance to each of the other species within the community is calculated, and the resulting values per focal species are averaged (mean pairwise distance per focal species, MPD_focal_; Box 1: Fig. 1). This simple extension enables quantification of the degree to which a given species co-occurs with other species, by measuring whether a nonnative species occurs more frequently with phylogenetically close/functionally similar or phylogenetically distant/functionally dissimilar species (Box 1: Fig. 1). The idea of focal-species is not novel *per se* and is conceptually similar to the α niche concept (Pickett and Bazzaz 1978), which can be interpreted as differences in functional traits between a given species and those of co-occurring species (Ackerly and Cornwell 2007). The focal-species approach as extended here is advantageous for incorporating different dimensions of biodiversity (i.e., taxonomic, functional, phylogenetic) and can be adapted at different spatial and temporal scales (e.g., Villalobos et al. 2013, 2016, Herrera-Alsina and Villegas-Patraca 2014, Miller et al. 2017a).

Considering previous studies that evaluated Darwin’s naturalization conundrum (e.g., Diez et al. 2008, Davies et al. 2011, Carboni et al. 2013, Bezeng et al. 2015, Marx et al. 2016), we expect (1) based on Darwin’s naturalization hypothesis, that nonnative species will tend to co-occur with distantly related species (overdispersion), or (2) based on the pre-adaptation hypothesis, that nonnative species will tend to co-occur more with closely related species (clustering) (Box1: Fig. 1C). The same logic extends to the influence of functional traits on species co-occurrence (Box 1: Fig. 1C): if co-occurring nonnative and native species are functionally similar, this would support the hypothesis that environmental fit mediates species co-occurrence within invaded communities; conversely, if co-occurring species are functionally distinct, this would support the premise that competition governs co-occurrence between native and nonnative species (Gallien et al. 2014, Carboni et al. 2018, Cadotte et al. 2018).

The fifty-year experimental fire frequency gradient at Cedar Creek (Peterson and Reich 2001, Reich et al. 2001, Willis et al. 2010, Cavender-Bares and Reich 2012), provides a unique opportunity to explore the changes in focal-species relatedness—in both phylogenetic and functional dimensions (Box 1: Fig. 1), given that nonnative species distribution and abundances are dynamic in time and space (Fig. S1) and the degree of interaction within invaded communities depends on species’ adaptability (Janzen 1985, Blackburn et al. 2015). The experiment thus provides a means to understand how stressful gradients influence focal-species phylogenetic relatedness/functional similarity, ergo Darwin’s naturalization conundrum. More specifically, as local conditions become more and/or differentially stressful (e.g., repeated fires versus deep shade at the two ends of the fire frequency gradient), species populations are expected to change, thereby increasing the role of species sorting (Nowacki and Abrans 2008, Cavender-Bares et al. 2009, Mayfield and Levine 2010, Willis et al. 2010, HilleRisLambers et al. 2012). Consequently, only species that have evolved to tolerate fire exposure and dry, high light conditions on the frequently burned end gradient, or to compete successfully for light at the unburned end of the gradient, are able to recruit and persist over time. Relevant functional traits—such as specific leaf area (cm^2^ g^-1^) [SLA], plant height (m), seed mass (mg) and rooting depth (m) that are related with dispersal facilitation, establishment, persistence, resource acquisition and recovery after disturbance (Moles et al. 2005, Peterson and Reich 2008, Willis et al. 2010, Cavender-Bares and Reich 2012, Díaz et al. 2015, Pinto-Ledezma et al. 2018)—are likely to change based on species niche requirements and resistance to disturbances.

Across fire gradients within the Cedar Creek system, plant height and tree cover are generally negatively associated with fire frequency, with taller and short plants distributed in low and high fire regimes, respectively; in turn, overstory tree cover decreases and light availability to the understory increases (Reich et al. 2001, Peterson and Reich 2008, Willis et al. 2010). SLA is predicted to decrease with increasing light availability in the oak savanna. In the understory of unburned dense forest canopies, high SLA maximizes light interception. In contrast, in frequently burned open areas prone to high solar radiation and desiccation, low SLA, which is associated with high leaf hydraulic resistance, limits plant desiccation (Givnish and Vermeij 1976, Ackerly 2004). Similarly, seed mass tends to decline with increasing fire frequency. Larger seeds are able to establish and survive as seedlings in competitive communities with low light availability, whereas small seeds are able to disperse farther and to reproduce in large numbers, allowing small-seed species to colonize patches that are not reached by large-seeded species (‘colonization-competition trade-off’ Tilman et al 1994; Leishman et al. 1995, Moles et al. 2005). Conversely, rooting depth tends to increase with increasing fire frequency, given that deeper roots enhance access to groundwater resources and increase plant recovery after fire disturbance (Peterson and Reich 2008, Willis et al. 2010, Pierret et al. 2016).

Considering species’ evolutionary as well as ecological differences and given that traits and phylogeny are linked due to underlying trait evolution (Box1: Fig. 1C), we test multiple predictions for explaining the variation in focal nonnative species phylogenetic relatedness/functional similarity across environmental gradients (Fig. 1). We posit that frequent fire—which is accompanied by nutrient depletion in this system—represents a higher degree of environmental stress than deep shade and heavy competition for light. Moreover, we hypothesize that there is antagonism between environmental stress and competitive interactions between the invader and its closely related native species—such that the likelihood of competitive exclusion weakens as environmental stress becomes greater. An alternative hypothesis would be that deep shade and strong light competition are equally stressful, but in other ways, than frequent fire. Testing our hypotheses, as articulated below, will illuminate these scenarios and how they relate to invasion and the DNC.

On the one hand, if species’ functional traits are conserved over evolutionary time (Box 1: Fig. 1C), we predict the relatedness of focal nonnative species to their native incumbents to shift from a pattern of phylogenetic overdispersion (low phylogenetic relatedness) to phylogenetic clustering (high phylogenetic relatedness) with increasing fire frequency (Fig. 1A). That is, we might expect that closely related and functionally similar plants—those with low SLA, deep roots and short height—will co-occur more in communities subjected to high fire frequency. We might further expect these same plants to be excluded from low frequency fire regimes that, instead, are dominated by functionally distinct and distantly related taller species that can persist under conditions of low understory light and soil fertility (e.g., shade-tolerant woody and herbaceous species; Cavender-Bares et al. 2004, Peterson and Reich 2008, Willis et al. 2010). On the other hand, if functional traits are convergent throughout species evolutionary history (Box 1: Fig. 1C), shifts from phylogenetic clustering to phylogenetic overdispersion are predicted with increasing fire frequency (Fig. 1B). That is, we might expect nonnatives to co-occurring more with distantly related but functionally similar natives with increasing fire frequency (Box 1: Fig. 1C). Notice that these predictions are based on the phylogenetic distance between nonnative species and their co-occurring species within the recipient communities (Box 1). Under this prediction, the steeper the environmental gradient—i.e., a gradient from unburned to high fire frequency (Fig. 1B)—the more nonnative species are able to avoid competition with ecologically similar native species (C_4_ grasses versus forbs; Leach and Givnish 1999) and as a consequence are not excluded from native communities subjected to high fire frequency where environmental filtering and sorting processes dominate due to the sunny, hot, dry, and nutrient-poor conditions where these communities are distributed (White 1983, Leach and Givnish 1999, MacArthur and Levins 1967, Peterson and Reich 2008, Cavender-Bares et al. 2009).

Real world possibilities may be complex. If biotic and abiotic filtering and sorting processes are acting simultaneously (Ackerly 2003, 2004, Swenson and Enquist 2009, Germain et al. 2018) and functional traits show phylogenetic signal—traits of the focal nonnative species could be either similar to (clustered) or distinct from (overdispersed) the native species in the community (Box 1: Fig. 1C); this would result in a complex nonlinear trend (polynomial) of changes in the focal nonnative species relatedness across the fire frequency gradient. For instance, as environmental conditions shift from communities with no fire to communities frequently subjected to fire (Fig. 1C), nonnative species would tend to co-occur mostly with closely related species, given that disturbances—such as fire regimes—filter or eliminate disturbance-sensitive species (Huston 1979). Thus, different fire regimes can select for fire-resistant (e.g., graminoids and forbs) species over shade-tolerant and fire-sensitive (e.g., woody plants) species (Peterson and Reich 2001, 2008). Once functionally similar species capable of tolerating conditions of frequent fire co-occur (e.g., grass-like plants and forbs), biotic forces may become more important such that nonnative species may co-occur with distantly related but ecologically similar (e.g., light-demanding and fire-resistant) species (Box 1: Fig. 1C).

## Methods

### Study site

A detailed description of the study site, burn units and sampling protocols can be found in Peterson and Reich (2001), Reich et al. (2001) and Cavender-Bares and Reich (2012) but is provided elsewhere.

The Cedar Creek Ecosystem Science Reserve is a 2300-ha reserve and National Science Foundation (NSF) Long Term Ecological Research since 1982 and is located on the Anoka Sand Plain in eastern Minnesota, USA. The climate is continental with cold and long winters and short, warm and humid summers. Mean annual temperature and mean annual precipitation are 6°C (ranging from ∼-12° in January and ∼22° in July) and ∼800 mm, respectively. The terrain is relatively flat, and soils varies according the elevation, with infertile, sandy and well drained in uplands areas and relatively fertile and poorly drained soils in lowland areas. Vegetation is variable from abandoned croplands to well preserved vegetation types, like dry oak savannas. The major vegetation types comprises deciduous forests, coniferous forests, deciduous woodlands and savannas, upland prairies and hardwood and coniferous swamps.

In order to characterize the responses of plant communities and to restore oak savanna vegetation a prescribed fire experiment started in 1964 (Peterson and Reich 2001, Cavender-Bares and Reich 2012). Within this fire experiment, an area of nearly 300 ha was divided into 19 management units ranging from 2.4 to 30 ha and each one was assigned a fire frequency treatment (five levels) that range from complete fire exclusion (unburned treatment) to yearly fire frequency (high fire frequency treatment). Following Cavender-Bares and Reich (2012) we categorized the fire frequency treatments into five levels: unburned (no fire), low frequency or Fire 1 (once per decade), medium frequency or Fire 2 (2-3*×* per decade), mid-high frequency or Fire 3 (4-5*×* per decade), and high frequency or Fire 4 (7-8*×* per decade). In addition, within each fire treatment, permanent sample plots of 0.375 ha (50 × 75 m) were established to collect plant species occurrence and abundance (Peterson and Reich 2008, Willis et al. 2010, Cavender-Bares and Reich 2012).

### General approach

Our aim was to evaluate both sides of Darwin’s naturalization conundrum (i.e., the DNH and PAH) by taking advantage of a long-term, frequently resampled (five-year intervals from 1984 to 2015) fire-frequency experiment that has established a gradient from unburned, dense woodland to frequently burned, open savanna. We evaluated DNH and PAH using taxonomic, functional, and phylogenetic information at local and landscape scales. At the local scale, species-level diversity metrics (see below) were estimated within each permanent plot of 0.375 ha (50 × 75 m) across fire experiments (*N* = 5) and for each five-year time interval (*N* = 7). For analysis at the landscape scale, species abundances and incidences were averaged across all plots representing each fire treatment by time interval combination.

### Functional and phylogenetic data

Data for four functional traits (specific leaf area [SLA], seed mass, plant height, and rooting depth)—selected based on the current understanding of key traits related to dispersal, establishment, resource acquisition, and persistence of plants across environmental gradients (Reich 2014, Moles 2017)—were obtained from previous studies (i.e., Willis et al. 2010, Cavender-Bares and Reich 2012). This trait dataset was not complete, with SLA available for 108 (44,44%) species, plant height for 86 (35,39%) species, and rooting depth for 85 (34,97%) species, respectively. We filled these gaps by supplementing the original trait dataset using the TRY (Kattge et al. 2011) and BIEN (Enquist et al. 2016) databases. In cases where missing traits were not available from these sources, we used genus-level averaged trait values.

The phylogenetic hypothesis was obtained from the recently published Spermatophyta mega-phylogeny (hereafter SB-tree; Smith and Brown 2018). The SB-tree was reconstructed under a hierarchical framework where individual major clades were first constructed and then placed into two different backbones (i.e, OTB [Open Tree of Life backbone] and MB [Magallón backbone]) and missing taxa were imputed—i.e., the insertion of missing species to an observed phylogeny—according to relationships in Open Tree of Life (see Smith and Brown 2018 for details). The final SB-tree contains a total of 353,185 and 356,305 species for the OTB and MB backbones, respectively, and to date is the most comprehensive phylogeny for seed plants (Smith and Brown 2018). In this study, we used the SB-tree constructed under the OTB backbone as this phylogenetic hypothesis provides more resolution towards the tips (Smith and Brown 2018). After checking species names in our community dataset and the pruned SB-tree OTB backbone, we found that 41 species were not sampled in the phylogenic tree, but these 41 species all had congeners represented in the SB-tree. Thus, to build a complete phylogenetic hypothesis, we added missing species into the pruned SB-tree using taxonomic constraints (i.e., adding terminal branches at the midpoint of their sister lineages) and estimated branching times under a birth-death model of diversification using the addTaxa package (Mast et al. 2015) in R version 3.4 (R Development Core Team 2018). We repeated the process 1000 times to account for phylogenetic uncertainty in the topology and the final phylogenetic hypothesis used in this study comprises a sample of 1000 fully dichotomous trees. Due to computational demand, we randomly selected a sample of 100 trees and all subsequent analyses were performed using these 100 trees.

### Nonnative species classification

We defined nonnative species as those that were introduced to the region by humans since early European-American activities in the region (∼1850s). Among the 243 vascular plants recorded in all permanent plots at the Cedar Creek Ecosystem Science Reserve, 26 species were identified as nonnative based on these criteria. Note that both nonnative and native plant species abundances are not stationary over time or across fire regimes, and certain plant species were not recorded in a particular time interval or fire experiment (Fig. S1).

### Calculation of diversity metrics

Taxonomic diversity was calculated by recording the number, abundance, and identity of species co-occurring with a nonnative focal-species in each permanent plot. We also estimated the pairwise species co-occurrence patterns by estimating the normalized checkerboard score (C-Score; Stone and Roberts 1990) between focal nonnative species and the co-occurring species within each plot (*N* = 5) in each fire treatment (*N* = 5) at each time interval (*N* = 7), i.e., 35 matrices per fire treatment for a total 175. This metric is commonly used to quantify species’ associations (Gotelli 2000, Bar-Massada 2015). We constructed null models that maintain species’ frequency and species richness within communities (i.e., permanent plots) (the trial-swap algorithm; Miklos and Podani 2004) to standardize C-Scores. This was done by calculating standardized effect sizes (SES), comparing an observed value (empirical C-Score) to the mean expected value under a null model, while accounting for variance (their standard deviation). For standardized C-Scores, positive and negative values denote segregated (negative species associations or competition) and aggregated (positive species associations or facilitation) patterns, respectively, while values close to zero, are consistent with random patterns (Callaway and Walker 1997, Tirado and Pugnaire 2005, Bar-Massada 2015).

Using functional and phylogenetic distance matrices as input data, we calculated different metrics accounting for abundance and/or presence-absence of co-occurring species with focal-species within each permanent plot. Prior to calculating functional metrics, a functional distance matrix was calculated using Euclidean distances given that our functional traits are continuous data. Metrics included mean pairwise phylogenetic and functional distances (MPD and MFD, respectively) and mean phylogenetic and functional nearest taxon distance (MPNTD and MFNTD, respectively). To facilitate comparisons and interpretation of the results, we also standardized phylogenetic and functional metrics using standardized effect sizes, with SES values >0 indicating phylogenetic or functional overdispersion and values <0 indicating clustering (Webb et al. 2002). The null model applied was the same used to standardize the C-Scores and the statistical significance of each metric was tested using thresholds (|SES| > 1.96) (Pinto-Ledezma et al. 2019). All calculations were conducted using customized scripts and functions modified from the picante (Kembell et al. 2010), metricTester (Miller et al. 2017b) and ecospat (Di Cola et al. 2017) packages in R.

Presence-absence and abundance-weighted metrics were highly correlated with one another (Fig. S2). Therefore, we only report results for the metrics abundance-weighted (aw) MPD/MFD and MPNTD/MFNTD because they are more sensitive for detecting sharp shifts in phylogenetic and functional structure (Miller et al. 2017b) and are more suitable for exploration of long-term data, such as ours.

### Phylogenetic signal in functional traits

We tested for phylogenetic signal in functional traits using Pagel’s *λ* (Pagel 1999) under a Bayesian approach. Pagel’s *λ* assumes a Brownian motion (BM) evolutionary model and its values range from 0 to 1, with values close to 0 and 1 indicating that traits evolved independently of the phylogeny (phylogenetic independence or no phylogenetic signal) and that traits evolved according to BM model (equivalent levels of phylogenetic covariance as expected under a BM model or phylogenetic signal), respectively (Münkemüller et al. 2012, Harmon 2018). We estimated Pagel’s *λ* for the same sample of 100 trees used in previous analyses and ran MCMC chains for 10 million generations, discarding the first million as burnins and sampling every 1000 generations in BayesTraits, version 3 (available from http://www.evolution.rdg.ac.uk/). Before this analysis, traits were log-transformed to meet the assumption of normality (Harmon 2018). We additionally estimated Pagel’s *λ* under a maximum likelihood approach and calculated values for Blomberg’s *K* (Blomberg et al. 2003). Both of these analyses produced results similar to those observed the Bayesian analysis of Pagel’s *λ*; we present only the latter for simplicity.

### Statistical analyses

To evaluate the probability of changes in the relatedness/similarity of focal-nonnative species to species in their surrounding communities through time (1984-2015) and across fire regimes (unburned to high-frequency), we modeled focal-species (for the three dimensions of diversity) as a function of fire regime (5 levels) and time interval (7 levels) under a Bayesian Multilevel Modeling (MLMs) framework (Gelman and Hill 2007, Finch et al. 2014). Using MLM models, metrics for each of the three dimensions of diversity were used to predict changes in focal-species relatedness by fire experiments and time periods—i.e., focal-species metrics values were nested within higher level units (fire experiments and time periods) (Finch et al. 2014). We used a bivariate tensor spline (Wood et al. 2013) to model the interaction effect of the predictors, which are of unknown, potentially non-linear form on focal-species relatedness (Bürkner 2017). MLMs were performed for all nonnative species together and for major taxonomic groups (i.e., dicots and monocots). In total, we constructed six models (3 levels x 2 metrics) related to changes in within-focal-species abundance and richness, 12 (3 levels x 4 metrics) related to changes in phylogenetic relatedness, and 12 (3 levels x 4 metrics) related to changes in functional similarity. In addition, given that environmental stressors like fire (or its absence) can influence species coexistence, we built several models in which species relatedness within fire experiments were a function of time intervals. This allowed us to explore how focal-species responded to environmental conditions over time. All MLMs were run using 4 NUTS sampling chains for 5000 generations, discarding 20% of each run as burnins, using the R package brms (Bürkner 2017), which implements Bayesian MLM in R using the probabilistic programming language Stan (Carpenter et al. 2017).

We also used permutational multivariate analysis of variance (PERMANOVA, Anderson 2001) to evaluate the uncertainty—the lack of complete knowledge about a parameter— associated with the imputation of missing species in the phylogenies. To do so, we modelled MPD/MPNTD as a function of fire regimes (5 levels) and time intervals (7 levels), using focal-species identities as random variables (N = 26) and phylogenetic trees (N = 100) as replicates. Under this modeling framework, the variance not explained by individual factors and their interactions can be attributed to differences resulting from phylogenetic uncertainty caused by species imputation (Rangel et al. 2015).

## Results

Native and nonnative species exhibited similar trait values to each other (Fig. S3). Statistical comparison—using Bayes factors (BF; Kass and Raftery 1995)—showed negligible evidence for differences in trait values between these two species categories (SLA [BF = 0.2725], plant height [BF = 0.2725], rooting depth [BF = 0.6065], and seed mass [BF = 0.2231]) (Fig. S4). Despite the similarity among natives and nonnatives, overall trait values varied considerably among all species at Cedar Creek (Fig. S4). In addition, while imputation of missing species in phylogenetic trees has been considered an important source of phylogenetic uncertainty in comparative studies, our results show that species imputation contributed little of the total variation in phylogenetic structure of focal-nonnative species (awMPD = 4.95%, wMPNTD = 10.10%). Most of the variation instead was attributable to evolutionary and ecological patterns of interest, i.e., phylogenetic and functional correlations among co-occurring species.

### Patterns of taxonomic, phylogenetic, and functional structure of focal-nonnatives

Species co-occurrence of focal nonnatives—measured as the tendency of focal species to occur in species-rich or species-poor communities—varied considerably among the 26 focal species (Fig. S5A-D). We found evidence for differences in species co-occurrence (abundance and richness) of focal nonnatives for dicots (Fig. S5 A and C; BF = 323.7592 and BF = 3498.187 for abundance and richness, respectively) but not for monocots (Fig. S5 B and D; BF = 0.3396 and BF = 0.4819 for abundance and richness, respectively). Co-occurrence patterns based on C-Scores also indicated high variability among focal species, although no statistical evidence was found for differences in species-pairs associations between dicots and monocots (Fig. S5E-F; BF = 0.2441 for dicots and BF = 0.1827 for monocots). Despite the lack of evidence for differences between species pairs, positive and negative interactions were observed between nonnatives and native species in recipient communities (Fig. S5E-F). For example, *Poa pratensis* (Kentucky bluegrass) that occurs in species rich and abundant communities (Fig. S5B-D), also tended to co-occur with species that interact positively with it (i.e., negative SES C-Score; Fig. S5-F).

In general, co-occurrence patterns for focal nonnative species varied strongly across both phylogenetic and functional dimensions (Fig. 2). All possible patterns were observed, e.g., greater co-occurrence with closely related but functionally distinct species (e.g., *Pyrus malus* [Fig. 2A and 2E] with *Digitaria sanguinalis* [Fig. 2B and 2F]), distantly related but functionally similar species (e.g., *Glechoma hederacea* [Fig. 2A and 2E] with *Setaria viridis* [Fig. 2B and 2F]), closely related and functionally similar species (e.g., *Linaria vulgaris* [Fig. 2A and 2E] with *Phleum pretense* [Fig. 2B and 2F]), and distantly related and functionally distinct species (e.g., *Plantago major* [Fig. 2A and 2E]). Despite this high degree of variation, there was no evidence for differences in co-occurrence patterns among dicots and among monocots (Fig. 2), whether for phylogenetic (BFs = 1.073 for dicots and 2.034 for monocots) or functional metrics (BFs = 0.270 for dicots and 2.034 for monocots).

**Fig. 2.**
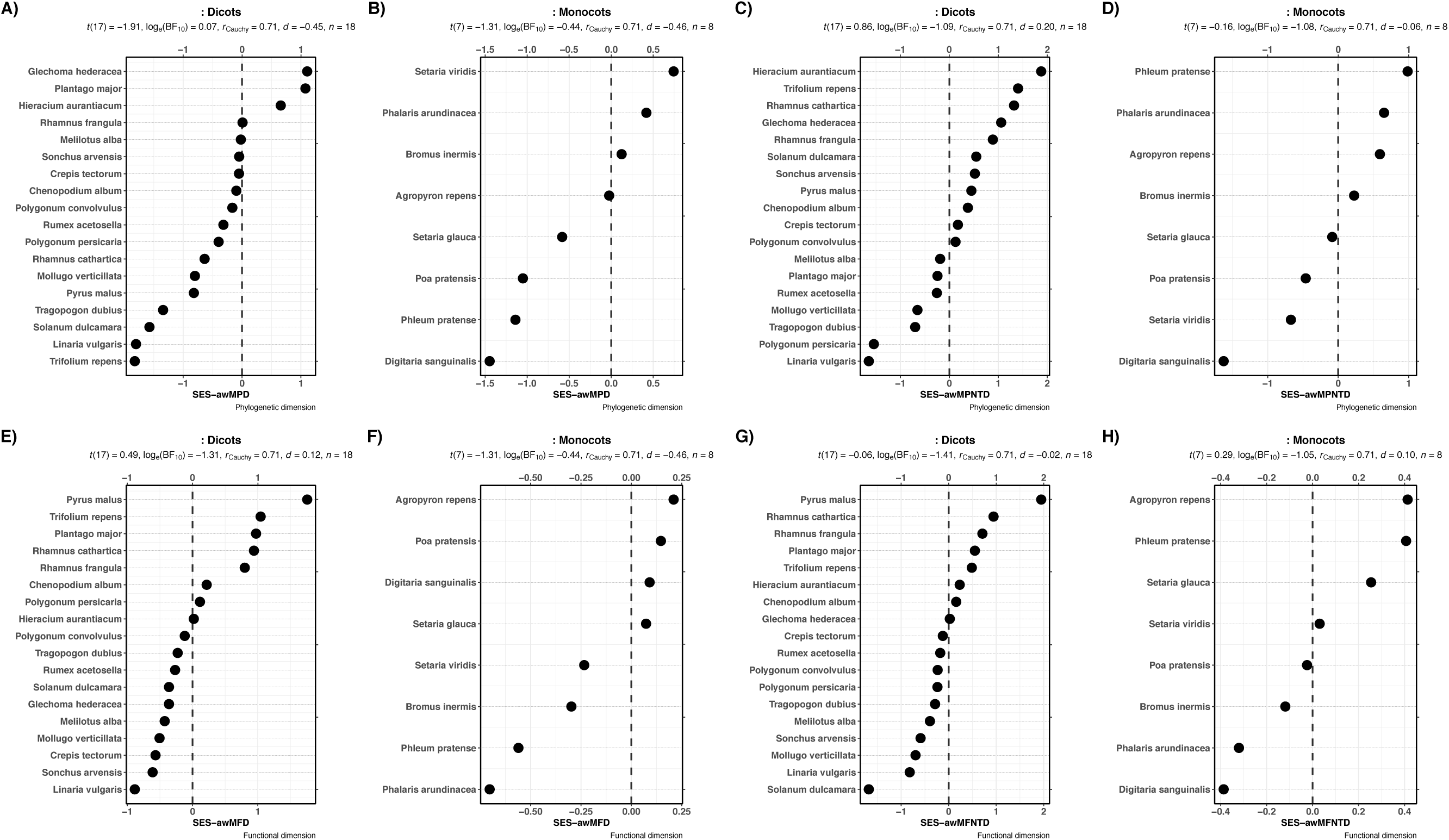
Patterns of phylogenetic relatedness and functional similarity of focal species relative to the plants in their surrounding communities for 26 nonnative species within the oak savanna understory permanent plots at the Cedar Creek Ecosystem Science Reserve, USA. Values correspond to abundance-weighted metrics averaged across all permanent plots (landscape scale) for both phylogenetic (top panel) and functional dimensions (bottom panel).

At the landscape scale, phylogenetic and functional metrics did not significantly differ from expectations under null models (Fig. 2), suggesting stochastic patterns of co-occurrence of nonnative species in recipient communities. However, at local scales we did detect significant deviations from null expectations (Fig. S6 and S7). For instance, the most common monocot and dicot species (*Poa pratensis* [Kentucky bluegrass] and *Polygonum convolvulus* [Black bindweed]) showed significant deviations from null expectations for phylogenetic clustering in 31% and 17% of all plots, respectively; and no significant deviations were found for phylogenetic overdispersion (Fig. S6). Additionally, in these same recipient communities, both Kentucky bluegrass and Black bindweed tended to co-occur more with functionally distinct species (overdispersion of traits, Fig. S7). Local-scale patterns showed shifts from overdispersion to clustering, and vice versa, across fire gradients and time periods for both phylogenetic and functional measures, indicating that co-occurrence patterns were dynamic over space and time across multiple axes of diversity (Fig. S6 and S7).

### Effects of fire regimes and time on co-occurrence patterns of focal nonnatives

For all three dimensions of diversity, the individual and combined effects of fire frequency and time were non-linear (Figs. 3 and 4). Focal nonnatives tended to co-occur more in species abundant (left panels in Fig. 3) and rich communities (middle panels in Fig. 3) with increasing fire frequency, although co-occurrence peaked at intermediate fire frequencies for some, e.g., abundance response for nonnative dicots (Fig. 3D). Co-occurrence under species-pairs associations (C-Score) showed complex, nonlinear patterns across fire experiments (right panels in Fig. 3), with positive associations found in the unburned and frequently burned treatments, and negative associations under intermediate fire frequencies. Notably, both dicot and monocot nonnatives responded similarly to fire gradients (Fig. 3, panels D to I).

**Fig. 3.**
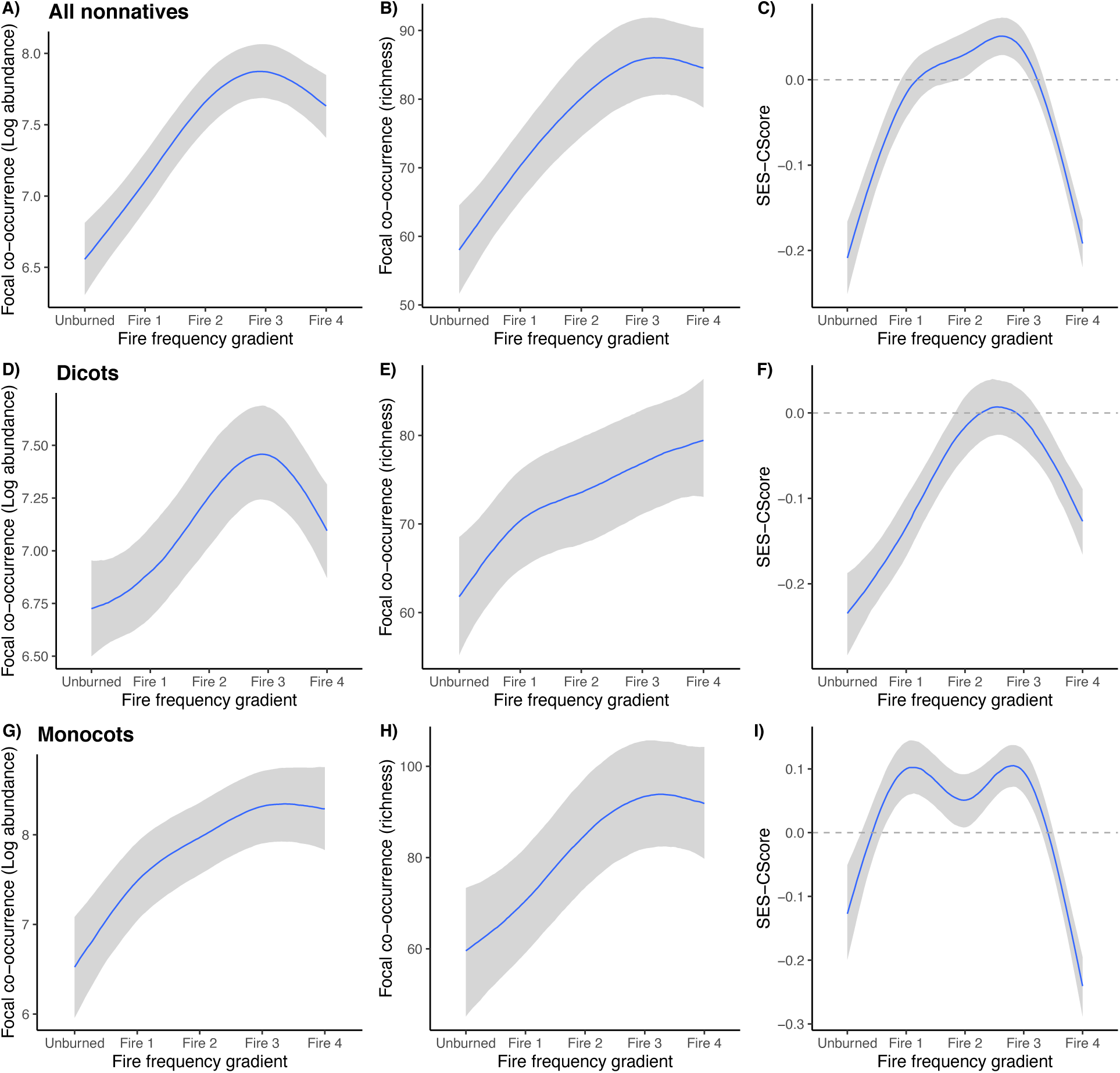
Marginal effect plots of changes in the taxonomic dimension across the fire gradient. Continuous blue lines represent the fitted slopes (with 95% confidence intervals in gray) that smooth over the fire frequency gradient from the Bayesian MLMs. Left-hand and middle columns show the variation of species co-occurrence of focal nonnatives for abundance and richness metrics, respectively and estimated as the tendency of focal nonnatives to occur in species-rich or species-poor communities. Right-hand column show the variation of focal species co-occurrence using the C-Score metric. Note that at the both extreme ends (i.e., unburned and fire 4 treatments) of the fire frequency gradient, nonnatives tend to co-occur with native species under positive interactions. Also, Bayesian MLMs show similar patterns for dicots and monocots, confirming that nonnative plant species respond in a similar way to fire regimes and not as a simple function of trees versus grass species.

**Fig. 4.**
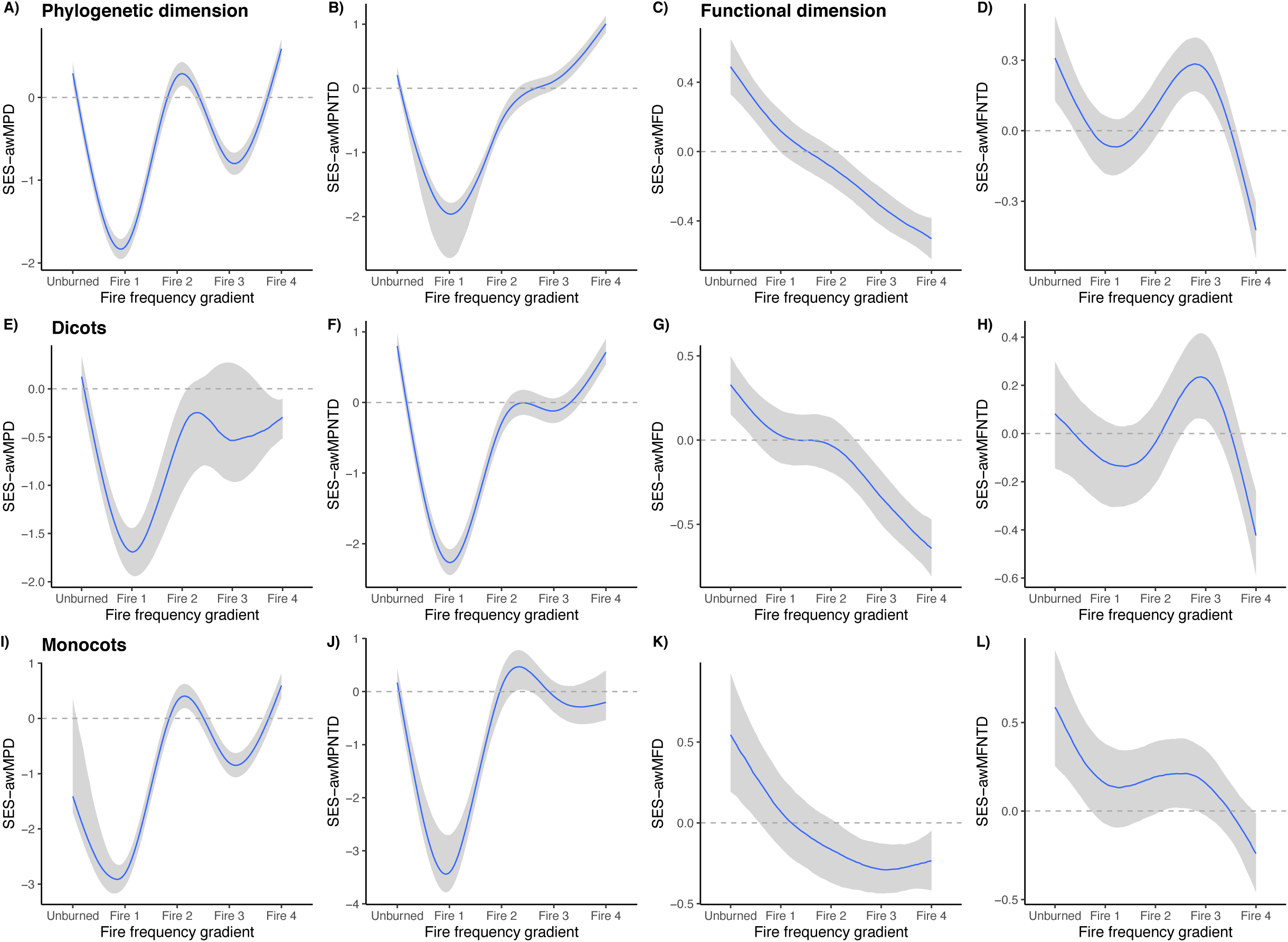
Marginal effect plots of changes in phylogenetic structure and functional structure of focal nonnative species in recipient communities across the fire frequency gradient. Continuous blue lines represent the fitted slopes (with 95% confidence intervals in gray) that smooth over the fire frequency gradient from the Bayesian MLMs. Left-hand columns (A, B, E, F, I and J) show the variation in focal nonnative species co-occurrence in recipient communities for the phylogenetic dimension and the right-hand columns (C, D, G, H, K and L) for the functional dimension. Y-axis for all panels represent fitted abundance-weighted metrics values (MPD/MFD and MPNTD/MFNTD) across the fire frequency gradient. See calculation diversity metrics in the method section for details of metric calculations.

Bayesian MLMs revealed a complex nonlinear response of focal nonnatives’ co-occurrence patterns to fire frequency (Fig. 4). These results were consistent regardless of phylogenetic scale—all species vs. dicots and monocots analyzed separately—and metric (awMPD/MFD, awMPNTD/MFNTD). Overall, there were changes from overdispersion to clustering in functional traits, with nonnative species tending to co-occur more with functionally similar species under increasing fire frequency (right-hand panels in Fig. 4). Furthermore, for the phylogenetic dimension, Bayesian MLM provided evidence for nonlinear changes in the phylogenetic structure of focal nonnatives in native communities with increasing fire frequency (left-hand panels in Fig. 4). Phylogenetic clustering was highest in communities subjected to low fire frequency, and phylogenetic overdispersion was greatest under intermediate and high-fire regimes, albeit with a return to phylogenetic clustering at medium-high fire regimes. Note that based on MPNTD, phylogenetic overdispersion was greatest under intermediate and high fire regimes (Fig. 4B).

Increased clustering of traits with increasing fire frequency was robust through time (Fig. 5 bottom panels). The phylogenetic dimension also revealed a complex temporal pattern (Fig 5. top panels). Although focal nonnatives tended to co-occur more with closely related species in unburned plots and with distantly related species in frequently burned plots, there were interannual changes in phylogenetic structure of nonnative species (Fig. 5). For example, there was a change from clustering to overdispersion in unburned plots in 2000 (Fig. 5 top-left panel), which may indicate that external factors (e.g., interannual climate variability) influenced co-occurrence patterns of focal nonnative species.

**Fig. 5.**
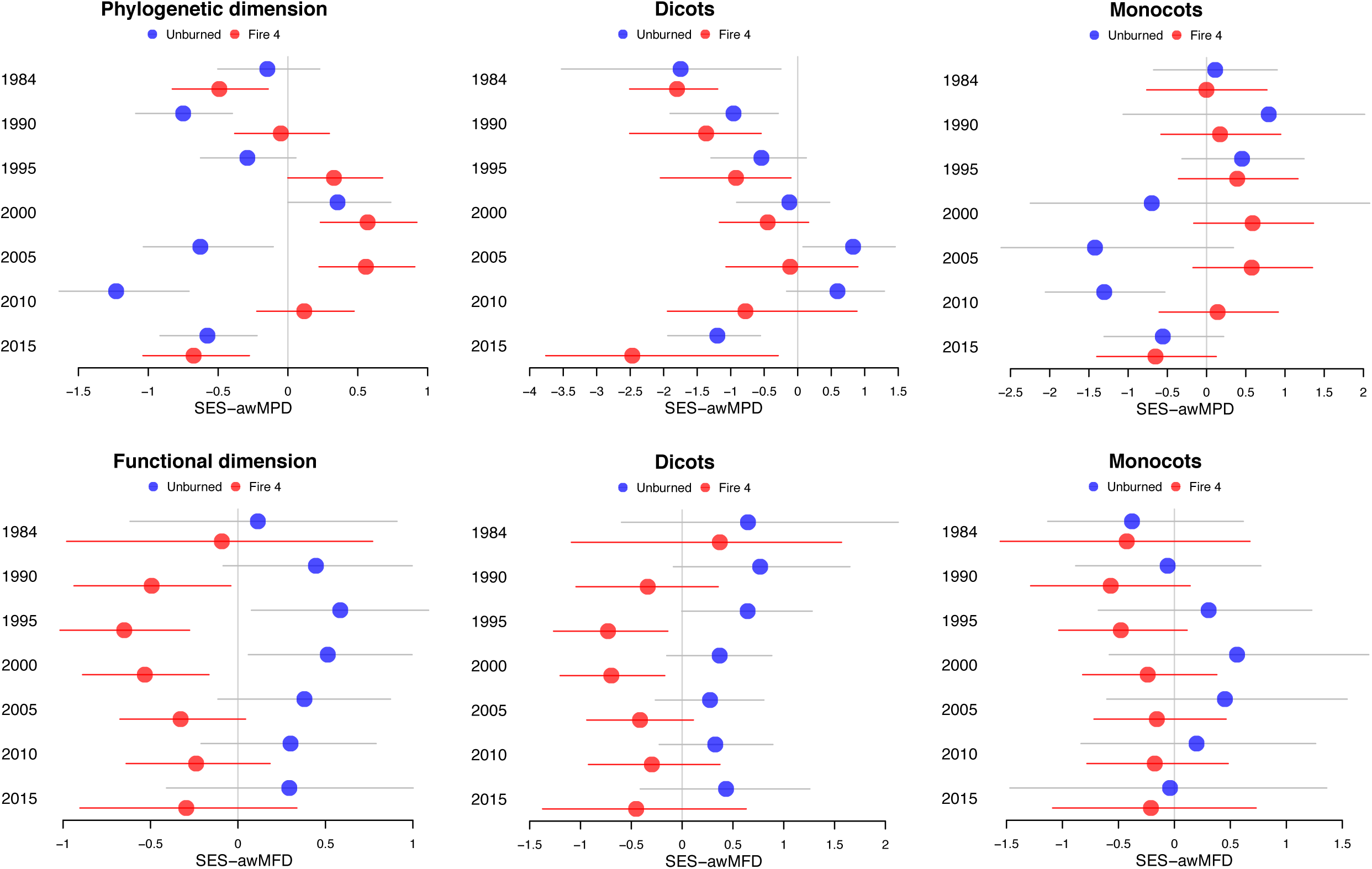
Fitted metric values from Bayesian multi-level models over time for the two extreme ends of the fire frequency gradient, i.e., unburned and fire 4 treatments. Upper and lower panels correspond to phylogenetic and functional dimensions, respectively.

### Phylogenetic signal in functional traits

Phylogenetic signal exhibited intermediate to high *λ* values (Fig. 6; though *λ* was weak for SLA), suggesting that trait values of co-occurring species in permanent plots at Cedar Creek resemble each other, although not at similar levels of phylogenetic covariance as would expected under a Brownian motion model of evolution (*λ* = 1; Fig. 6). These results were consistent across all 100 phylogenetic trees, i.e., regardless of phylogenetic uncertainty introduced by missing terminal branches.

**Fig. 6.**
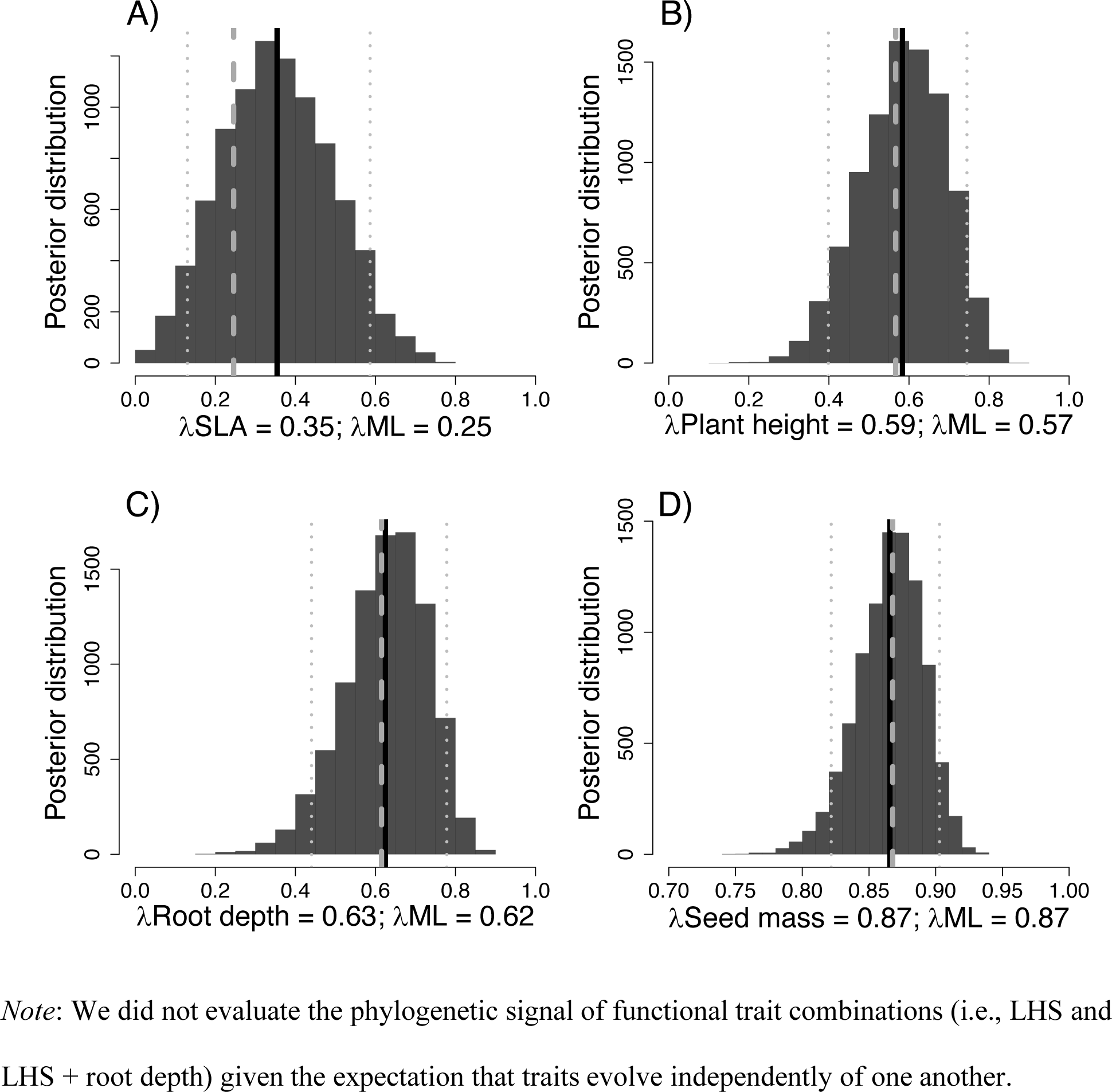
Bayesian phylogenetic signal estimated using Pagel’s *λ* and a sample of 100 phylogenetic trees. Posterior probability distributions of (A) specific leaf area, (B) plant height, (C) rooting depth, and (D) seed mass. Posterior distribution (and 95% confidence interval, vertical gray dotted lines) show *λ* values from 10 million generations sampled every 1000 generations. Vertical dashed gray line represent the mean *λ* value estimated under a maximum likelihood approach over a sample of 100 trees.

## Discussion

The temporal dynamics of co-occurrence patterns of nonnative species in recipient oak savanna communities across a long-term experimental fire gradient revealed that neither of Darwin’s competing hypotheses fully explain species invasiveness in plant community assembly. We found that co-occurrence of the 26 nonnative species within invaded communities (Fig. S1) did not follow a general tendency of clustering (which would support the pre-adaptation hypothesis) or overdispersion (which would support Darwin’s naturalization hypothesis) (Fig. 2). Instead, we found that community assembly was influence by species differences in evolutionary history (i.e., shared ancestry) and by ecological differences associated with functional traits that ultimately regulate the colonization and persistence of nonnatives within invaded communities (Box 1: Fig. 1C).

Applying our focal-species approach to three different dimensions of vascular plants diversity (i.e., taxonomic, phylogenetic and functional) in communities sampled over decades across an experimental fire gradient, we found that nonnatives tend to be phylogenetically distantly related to the recipient communities in the unburned treatment and in the high fire frequency treatment. In other words, nonnatives are closely related to native species in recipient communities—except in the extreme ends of the fire gradient, where they are more distantly related than expect—which might suggest a filtering process at intermediate fire frequencies favoring the co-occurrence of phylogenetically close relatives. This pattern is consistent between monocots and dicots and contrast with functional similarity. Indeed, nonnatives are functionally dissimilar in the unburned treatment, consistent with biotic interactions that limit ecologically similar species from co-occurring. As fire frequency increases, nonnatives species become increasingly similar in function to the recipient native species, indicating that similar functions are required for the persistence of nonnative species in recipient communities subjected to strong environmental filtering caused by the fire regimes. Importantly, these patterns are similar within both monocots and dicots; hence not simply function of tree versus grass dynamics. Moreover, when co-occurrence of nonnatives to recipient natives was evaluated through time, new insights regarding the assembly processes emerge. We found a consistent pattern through time for the functional dimension, with nonnatives co-occurring with functionally dissimilar and similar natives in the unburned and the highest fire regimen treatments, respectively. The phylogenetic dimension, on the other hand, showed shifts in nonnative co-occurrence patterns that suggest additional external forces influencing the phylogenetic relatedness of nonnatives with their recipient native species. Altogether, our findings provide insights for understanding the invasion dynamics of native communities across environmental gradients and at multiple temporal and spatial scales in a highly diverse region of North America.

The degree to which (and the mechanisms at work by which) nonnative species successfully invade new communities has remained controversial (Prins and Gordon 2014), particularly because a variety of assembly processes are involved in the effective establishment of introduced species outside their native ranges (Gallien and Carboni 2017, Cadotte et al. 2018, Redding et al. 2019). This has generated growing interest in the role of evolutionary and ecological differences among taxa in predicting nonnative species invasiveness (Carboni et al. 2013, Gallien and Carboni 2017, Cadotte et al. 2018). Studies have reported contrasting results in terms of phylogenetic relatedness of nonnative species to recipient assemblages (Carboni et al. 2013, Li et al. 2015, Marx et al. 2016), with mixed support for both hypotheses comprising Darwin’s naturalization conundrum, i.e., Darwin’s naturalization hypothesis and pre-adaptation hypothesis (Kusumuto et al. 2019). Our results showed differential patterns of relatedness of focal nonnative species to recipient native communities, with some nonnatives co-occurring more with closely related natives, and others with distantly related natives (Fig. 2 upper panel). These results suggest that different assembly processes—e.g., biotic interactions, environmental filtering—have simultaneous effects on focal nonnative co-occurrence patterns (Boulangeat et al. 2012, Gallien and Carboni 2017, Kusumuto et al. 2019; see also Box 1: Fig. 1C). Integrating multiple dimensions of diversity (i.e., taxonomic, phylogenetic, and functional) in the study of invasions can offer new insights about assembly processes driving nonnative species co-occurrence patterns (Gallien and Carboni 2017, Cadotte et al. 2018).

Evaluating the taxonomic dimension, we found that both positive and negative interactions appear to be important in the co-occurrence of nonnative species with native species in invaded communities (Fig. S5E-F). Functional patterns differed from those observed for the phylogenetic dimension (Fig. 2 lower panel). This may indicate that functional traits are modulating co-occurrence patterns between nonnative and native species (Marx et al. 2016, Carboni et al. 2018), by controlling or regulating the species response to different assembly processes (Cavender-Bares et al. 2004, Swenson and Enquist 2009, Pinto-Ledezma et al. 2018). For example, similarity in ecologically relevant traits of closely related species should result in patterns of phylogenetic clustering under similar environmental conditions, while similarity in distantly related species would be associated with a pattern of phylogenetic overdispersion (Cavender-Bares et al. 2004, 2009). In other words, evolutionary conservatism and convergence of functional traits help to explain why nonnative species tend to co-occur with closely related natives (as suggested in PAH) and distantly related natives (as indicated by DNH), respectively (Box 1: Fig. 1C) because functional traits in co-occurring species at Cedar Creek resemble each other. In other words, nonnative species have similar trait values to those of native species (Fig. a pattern that is expected to emerge over time due to environmental sorting and not by random introductions of species in the study region.

Species composition and structure within communities is dynamic in space and time, varying from short timeframes at small scales (e.g., Fig. S1) to millions of years across biogeographical regions (Chesson 2000, Williams et al. 2004, Cavender-Bares and Reich 2012, Pinto-Ledezma et al. 2018). Thus, a better understanding of invasion dynamics and their relationship to DNC requires evaluation across multiple temporal scales. Indeed, recent evidence indicates that a single snapshot in time could mislead interpretations on both sides of Darwin’s conundrum (Li et al. 2015, Cadotte et al. 2018) or prevent observation of the full range of impacts of nonnative species on invaded communities (Gilbert and Levine 2013). Our results support these time-scale dependent findings, as we found shifts over time in the phylogenetic and functional co-occurrence structure of focal nonnatives across the fire frequency gradient (Fig. S6 and S7). For example, considering a single time period, *Poa pratensis* (Kentucky bluegrass)— the most common and abundant nonnative species in our study region (Fig. S1)—tended to co-occur more with closely related species, supporting the PAH (Fig. 2B); however, in plots protected from fire, phylogenetic clustering from 1984 to 2005 gave way to phylogenetic overdispersion in 2010 and 2015 (Fig. S6). Thus, depending on the time periods considered, the co-occurrence patterns of *Poa pratensis* could support PAH or DNH. These findings highlight the importance of multi-temporal evaluations in the study of biological invasions, given that co-occurrence patterns are an emergent characteristic of dynamic population changes across many species (Chesson 2000, Cavender-Bares and Reich 2012, Li et al. 2015). Indeed, an observed species may be only fleetingly present in a community, because it recently colonized it (Gilbert and Levine 2013, Germain et al. 2018).

Environmental conditions and disturbances constrain patterns of species diversity within communities by acting as filters that alter species co-occurrence patterns (Connell 1978, Peterson and Reich 2008, Pinto-Ledezma et al. 2018). Disturbances such as fire can generate stressful environmental conditions in which only a subset of tolerant species will recruit and persist, thereby diminishing the strength of density-dependency interactions (Peterson and Reich 2008, HilleRisLambers et al. 2012, Coyle et al. 2014). These factors also influence the degree to which nonnative species co-occur with natives in invaded communities (Thuiller et al. 2010, Cadotte et al. 2018). Our results show that fire frequency generated idiosyncratic effects on co-occurrence patterns in invaded communities (Fig. 4), indicating the simultaneous and interacting effect of assembly processes across environmental gradients—i.e., as environmental conditions change, density-dependent interactions promote the coexistence of some species while excluding others (Germain et al. 2018). In fact, we found that functional similarity of focal nonnatives to natives increased with increasing fire frequency, which might suggest a strong influence of environmental filtering (right panels in Fig. 4). However, phylogenetic distance exhibited a complex nonlinear trend (left panels in Fig. 4), matching our third prediction (Fig. 1C). Specifically, phylogenetic clustering of nonnatives to natives was maximized under low fire frequency. Shifts to phylogenetic overdispersion appeared under medium-high fire frequency, possibly as a consequence of competitive interactions (Fig. 4). In other words, both environmental filtering and biotic interactions are acting in concert in shaping co-occurrence patterns of nonnative species across environmental gradients. This is consistent with prior experimental studies (e.g., Germain et al. 2018) highlighting the importance of simultaneous effects of environmental filtering and competitive interactions on community assembly (Chesson 2000, Ackerly 2003, Germain et al. 2018).

### Perspectives

Extending the diversity fields’ framework to focal-species under combined phylogenetic and functional perspective (Box 1) offers an innovative way to evaluate patterns of species-level co-occurrence at local scales, enhancing our understanding of the mechanisms driving the assembly of natural communities. For example, by combining different dimensions of vascular plant diversity under the focal-species framework, we were able to detect shifts in the phylogenetic relatedness and functional similarity of focal nonnatives with respect to recipient communities across environmental gradients and time periods. This is particularly important given that individual species—including nonnative species—respond differently to changes in the environmental conditions of local communities (Tilman 2004, Ackerly and Cornwell 2007, Li et al. 2015, Redding et al. 2019). Further analyses addressing different taxa, biogeographical regions, or environmental conditions could reveal more insights about the mechanisms driving co-occurrence of nonnative species in recipient communities, or elucidate the role of other axes of variation shaping species co-occurrence patterns. Finally, the integration of multiple dimensions of biodiversity within the “focal species” framework can enhance our ability to produce reliable information of species co-occurrence at different spatial and temporal scales, facilitating our ability to monitor changes in both individual species and whole communities, and thus helping to guide conservation efforts.

## Acknowledgments

J.N.P-L. was supported by the University of Minnesota College of Biological Sciences’ Grand Challenges in Biology Postdoctoral Program. Data collection and archiving and maintenance of the fire frequency experiment were supported by the Cedar Creek NSF Long-Term Ecological Research program (DEB-1234162; DEB-1831944). FV was supported by CONACYT through INECOL, Mexico.

## Statement of authorship

JC-B, JNP-L and DJL conceived the ideas presented and tested herein. JNP-L managed the project. JNP-L performed the analyses, and wrote the first draft. FV and PBR contributed to the ideas and interpretation of data, and all authors contributed throughout the whole writing process. PBR has co-led the implementation and management of the long-term fire frequency experiment and associated community censusing and data curation.

## BOX 1

Species co-occurrence patterns in local communities are a consequence of ecological processes (species interactions with other organisms and the environment) and historical processes (biogeographic history, long-term dispersal, past diversification) that operate over short and long time scales, respectively (Cavender-Bares et al. 2016, 2018, Pinto-Ledezma et al. 2019). Different metrics have been used to measure the degree of relatedness/similarity (see Miller et al. 2017, Scheiner et al. 2017) of species co-occurring within local communities to infer assembly processes from community phylogenetic/functional structure (Cavender-bares et al. 2006, Mayfield and Levine 2010). These metrics are calculated using a community data matrix (CDM)—in which rows are communities and columns are species—in combination with a distance matrix, either phylogenetic or functional. Generally, CDMs are analyzed by rows (*Q*-mode in Arita et al. 2008, see also Villalobos and Arita 2010), summarizing information at the community level; however, CDMs can also be analyzed by columns (*R*-mode in Arita et al. 2008, Villalobos and Arita 2010), summarizing information at the species level. In addition, by intersecting rows and columns (*Qr*-mode and *Rq*-mode in Arita et al. 2008), it is possible to obtain new information, e.g., the ‘diversity field’ of a given species (Arita et al. 2008), which represents the species richness of communities within the distribution of a species. This can be interpreted as the tendency of a given species to occur in species-rich or species-poor communities, i.e., to coexist with many or with few species (Villalobos and Arita 2010).

Building on this framework, Villalobos et al. (2013, 2017) extended the concept of diversity fields—the set of diversity values of sites within the geographical range of a given species (Arita et al. 2008)—to ‘phylogenetic fields’ (Villalobos et al. 2013) and ‘functional fields’ (Villalobos et al. 2017, see also Miller et al. 2017) that describe the overall phylogenetic/functional structure of species co-occurring with a given species geographical range. These fields are interpreted as the tendency of a focal species to co-occur either with closely related/functionally similar or distantly related/functionally dissimilar species (Villalobos et al. 2013, 2017). Similarly, phylogenetic/functional fields can be used as metrics of species-level coexistence. All three of these approaches—diversity, phylogenetic, and functional fields— are usually applied at macroecological scales (Arita et al. 2008, 2010, Villalobos and Arita 2010, Villalobos et al. 2013, 2016, 2017) with some local-scale exceptions (e.g., Elliot et al. 2016, Miller et al. 2017a, Kusumuto et al. 2019). Given that the observational units in our study are species within local communities, we downscale the concept of fields applied at large geographical scales to co-occurring species within local communities (Box1: Fig. 1A-B). This extension enables evaluation of co-occurrence patterns and inferences regarding assembly processes to be applied at the level of individual species rather than entire communities. In doing so, we obtain separate estimates for each species occurring in a local community rather than a single mean across all species co-occurring in a community (Box 1: Fig. 1A-B).

**Box 1: Fig. 1.**
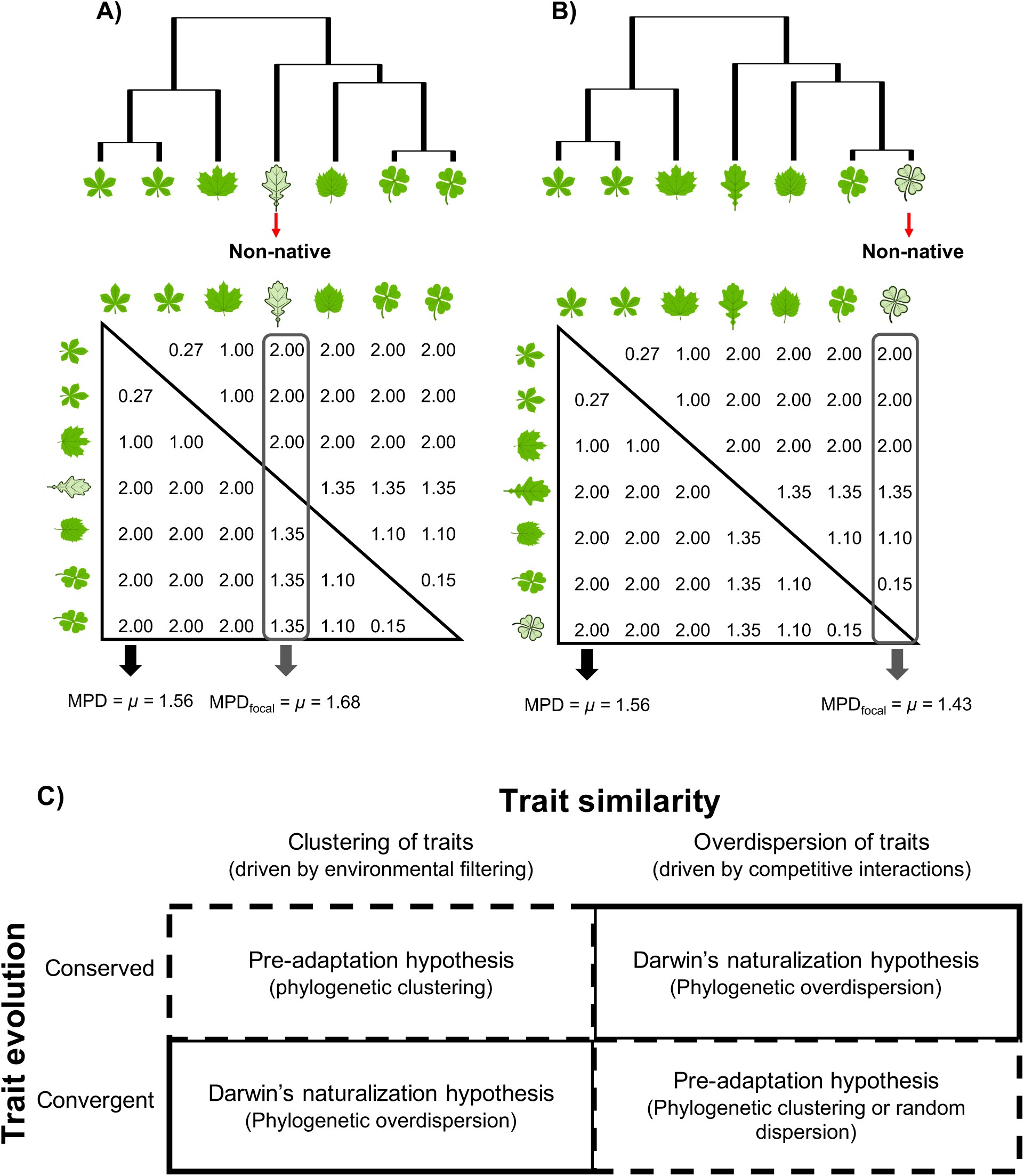
Conceptual framework for the estimation phylogenetic/functional structure of focal-species contrasted with the traditional estimation of phylogenetic/functional structure at the community level. Here we show the difference between estimates for standard mean pairwise phylogenetic distance (MPD) between species occurring within a community (a single, community-level value) and MPD_focal_ (multiple, species-level values). Again, while traditional MPD is calculated based on the mean distance among all possible pairs of species within a community, MPD_focal_ is based on the mean distance of a focal species relative to the co-occurring species in a local community (e.g., nonnative species in A and B). Under this simple framework, it is possible to evaluate both the phylogenetic relatedness and functional similarity of nonnative species in native communities and consequently conduct a more comprehensive analysis of both sides of Darwin’s naturalization conundrum (C). (A) Darwin’s naturalization hypothesis (DNH)—that nonnative species closely related to native species are less likely to successfully invade native communities because they share similar and already occupied niches (continuous boxes in panel C). (B) Pre-adaptation hypothesis (PAH)—that nonnative species closely related to native species should be favored because they share similar traits with native species, making them well-suited (“preadapted”) to the novel range, and permitting them to colonize and further adapt (dashed boxes in panel C). Panel C adapted from Cavender-Bares et al. (2004)

