## Supplementary figures for "Testing Darwin’s naturalization conundrum based on taxonomic, phylogenetic and functional dimensions of vascular plants"

**Fig. S1.** Relative abundance of nonnative species in the oak savanna understory permanent plots at Cedar Creek Ecosystem Science Reserve, Minnesota, USA. Colors depict the fire treatment and circle sizes are proportional to the relative abundance (log transformed) averaged by fire treatment. Species names are colored according the major groups of vascular plants; monocots = orange and dicots = green.

**Fig. S2.** Correlation matrices for the metrics of phylogenetic and functional dimensions evaluated in this study.

**Fig. S3.** Maximum clade credibility tree obtained from 1000 trees and the distribution of the four functional traits (log transformed for graphical representation) used in this study. Species names in red correspond to nonnative species.

**Fig. S4.** Comparison of functional traits between native and nonnative species. A) SLA, B) plant height, C) rooting depth, and d) seed mass.

**Fig. S5**. Patterns of co-occurrence for the taxonomic dimension over time and across the fire frequency gradient.

**Fig. S6.** Patterns of phylogenetic structure of focal nonnative species over time and across the experimental fire gradient.

**Fig. S7.** Patterns of functional structure of focal nonnative species over time and across the experimental fire gradient .

**Fig. S1**

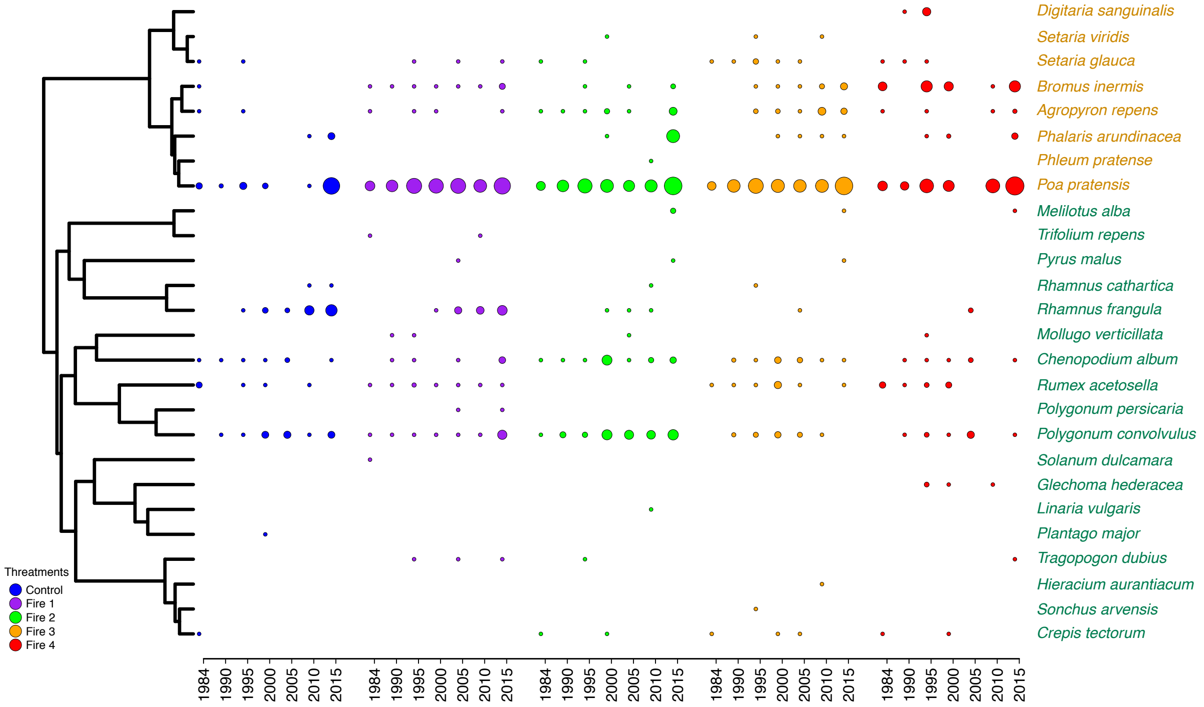

Digitaria sanguinalis
Setaria viridis
Setaria glauca
Bromus inermis
Agropyron repens
Phalaris arundinacea
Phleum pratense
Poa pratensis
Melilotus alba
Trifolium repens
Pyrus malus
Rhamnus cathartica
Rhamnus frangula
Mollugo verticillata
Chenopodium album
Rumex acetosella
Polygonum persicaria
Polygonum convolvulus
Solanum dulcamara
Glechoma hederacea
Linaria vulgaris
Plantago major
Tragopogon dubius
Hieracium aurantiacum
Sonchus arvensis
Crepis tectorum
Threatments
Control
Fire 1
Fire 2
Fire 3
Fire 4
1984 1990 1995 2000 2005 2010 2015
1984 1990 1995 2000 2005 2010 2015
1984 1990 1995 2000 2005 2010 2015
1984 1990 1995 2000 2005 2010 2015
1984 1990 1995 2000 2005 2010 2015

**Fig. S2**

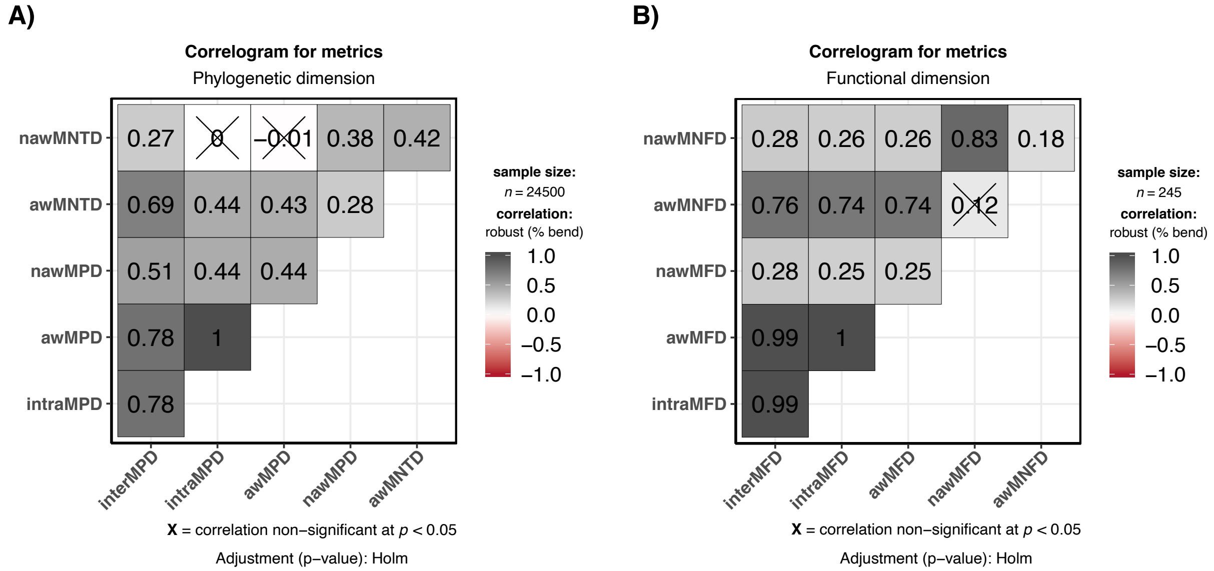
A)
Correlogram for metrics
Phylogenetic dimension
nawMNTD
awMNTD
nawMPD
awMPD
intraMPD
0.27
0
−0.01
0.38
0.42
0.69
0.44
0.43
0.28
0.51
0.44
0.44
0.78
1
0.78
interMPD
intraMPD
awMPD
nawMPD
awMNTD
sample size:
n = 24500
correlation:
robust (% bend)
1.0
0.5
0.0
−0.5
−1.0
X = correlation non−significant at p < 0.05
Adjustment (p−value): Holm
B)
Correlogram for metrics
Functional dimension
nawMNFD
awMNFD
nawMFD
awMFD
intraMFD
0.28
0.26
0.26
0.83
0.18
0.76
0.74
0.74
0.12
0.28
0.25
0.25
0.99
1
0.99
interMFD
intraMFD
awMFD
nawMFD
awMNFD
sample size:
n = 245
correlation:
robust (% bend)
1.0
0.5
0.0
−0.5
−1.0
X = correlation non−significant at p < 0.05
Adjustment (p−value): Holm

**Fig. S3**

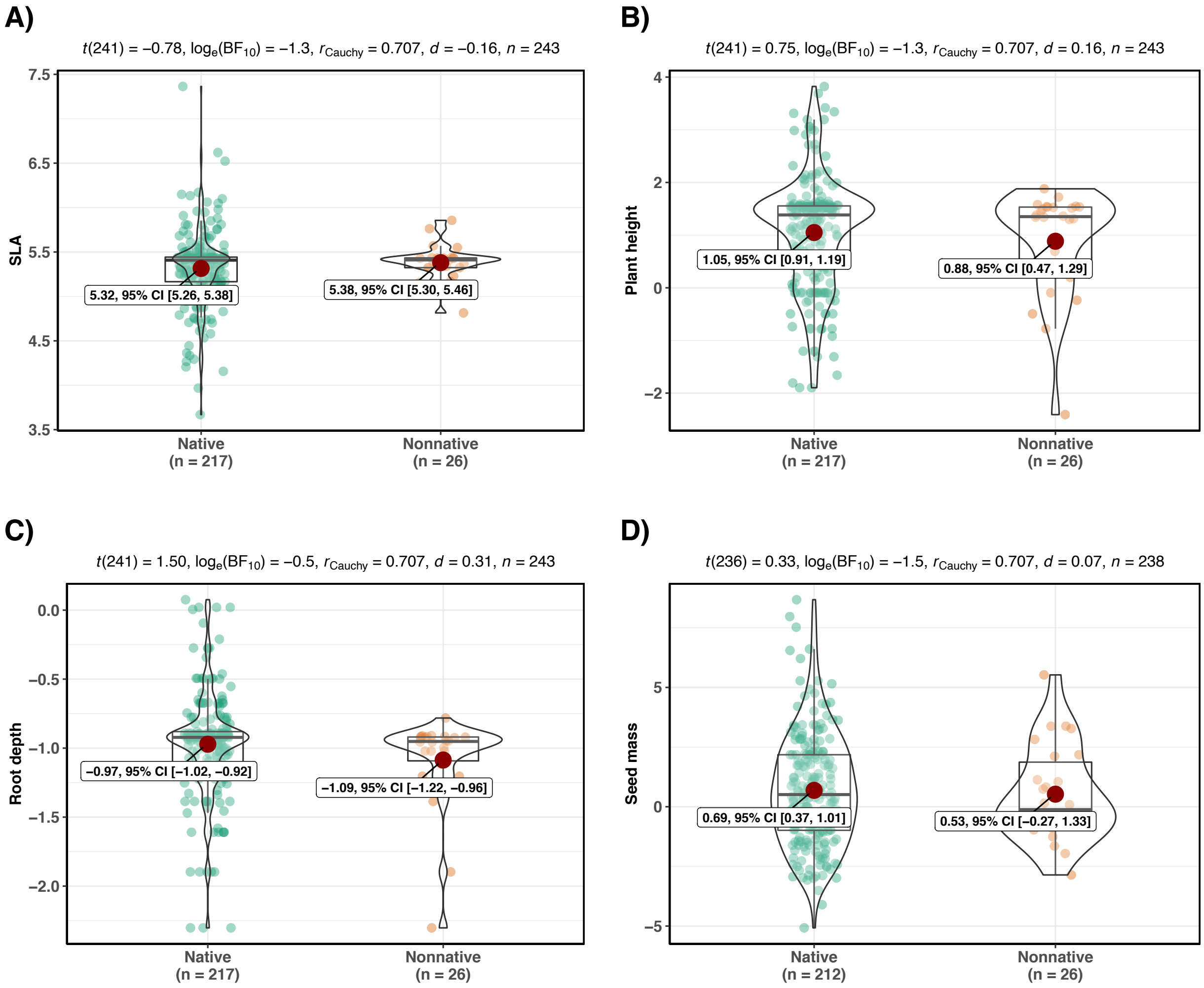
A)
t(241) = −0.78, loge(BF10) = −1.3, rCauchy = 0.707, d = −0.16, n = 243
SLA
7.5
6.5
5.5
4.5
3.5
5.32, 95% CI [5.26, 5.38]
5.38, 95% CI [5.30, 5.46]
Native
(n = 217)
Nonnative
(n = 26)
B)
t(241) = 0.75, loge(BF10) = −1.3, rCauchy = 0.707, d = 0.16, n = 243
Plant height
4
2
0
−2
1.05, 95% CI [0.91, 1.19]
0.88, 95% CI [0.47, 1.29]
Native
(n = 217)
Nonnative
(n = 26)
C)
t(241) = 1.50, loge(BF10) = −0.5, rCauchy = 0.707, d = 0.31, n = 243
Root depth
0.0
−0.5
−1.0
−1.5
−2.0
−0.97, 95% CI [−1.02, −0.92]
−1.09, 95% CI [−1.22, −0.96]
Native
(n = 217)
Nonnative
(n = 26)
D)
t(236) = 0.33, loge(BF10) = −1.5, rCauchy = 0.707, d = 0.07, n = 238
Seed mass
5
0
−5
0.69, 95% CI [0.37, 1.01]
0.53, 95% CI [−0.27, 1.33]
Native
(n = 212)
Nonnative
(n = 26)

**Fig. S4**

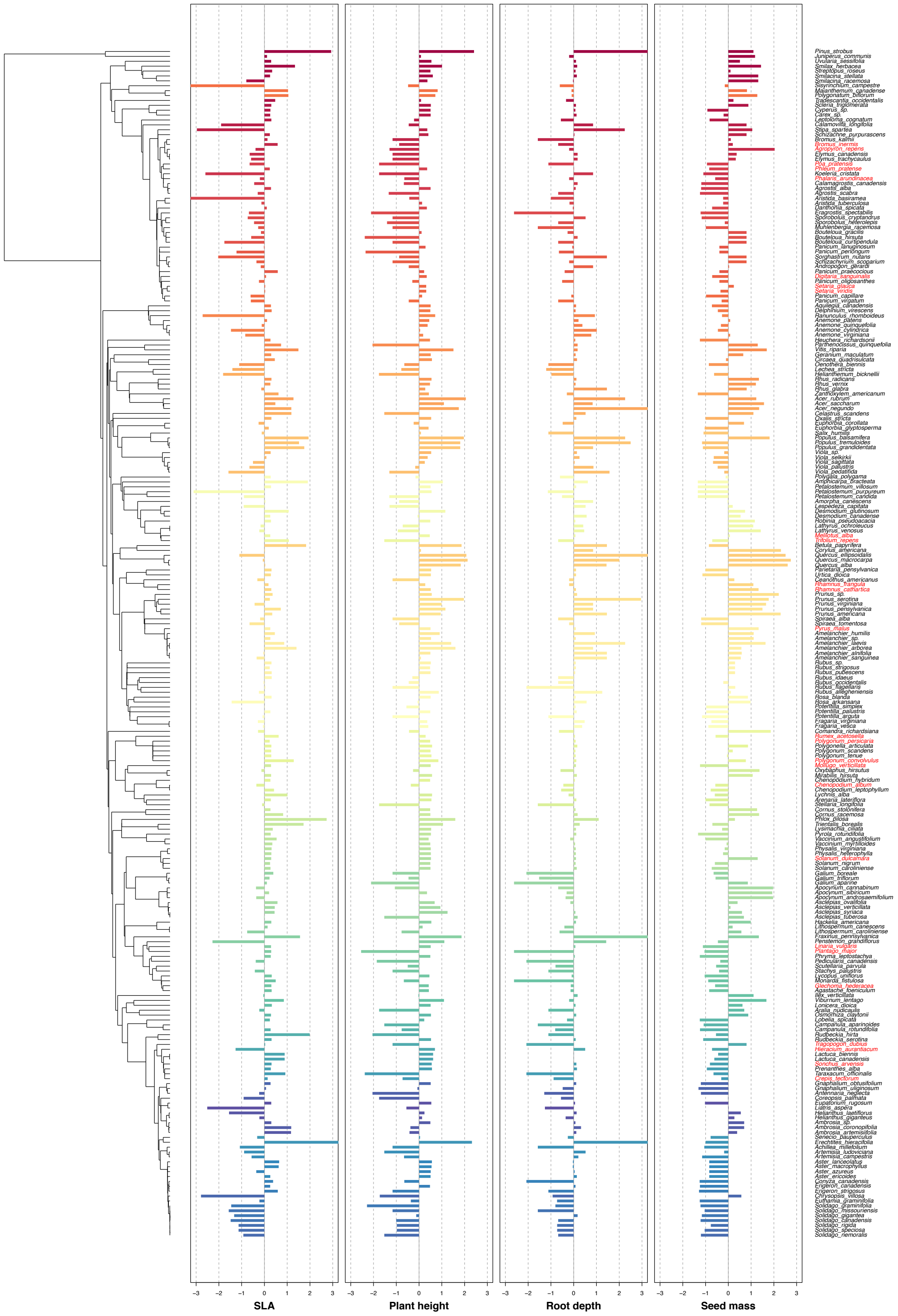

**Fig. S5**

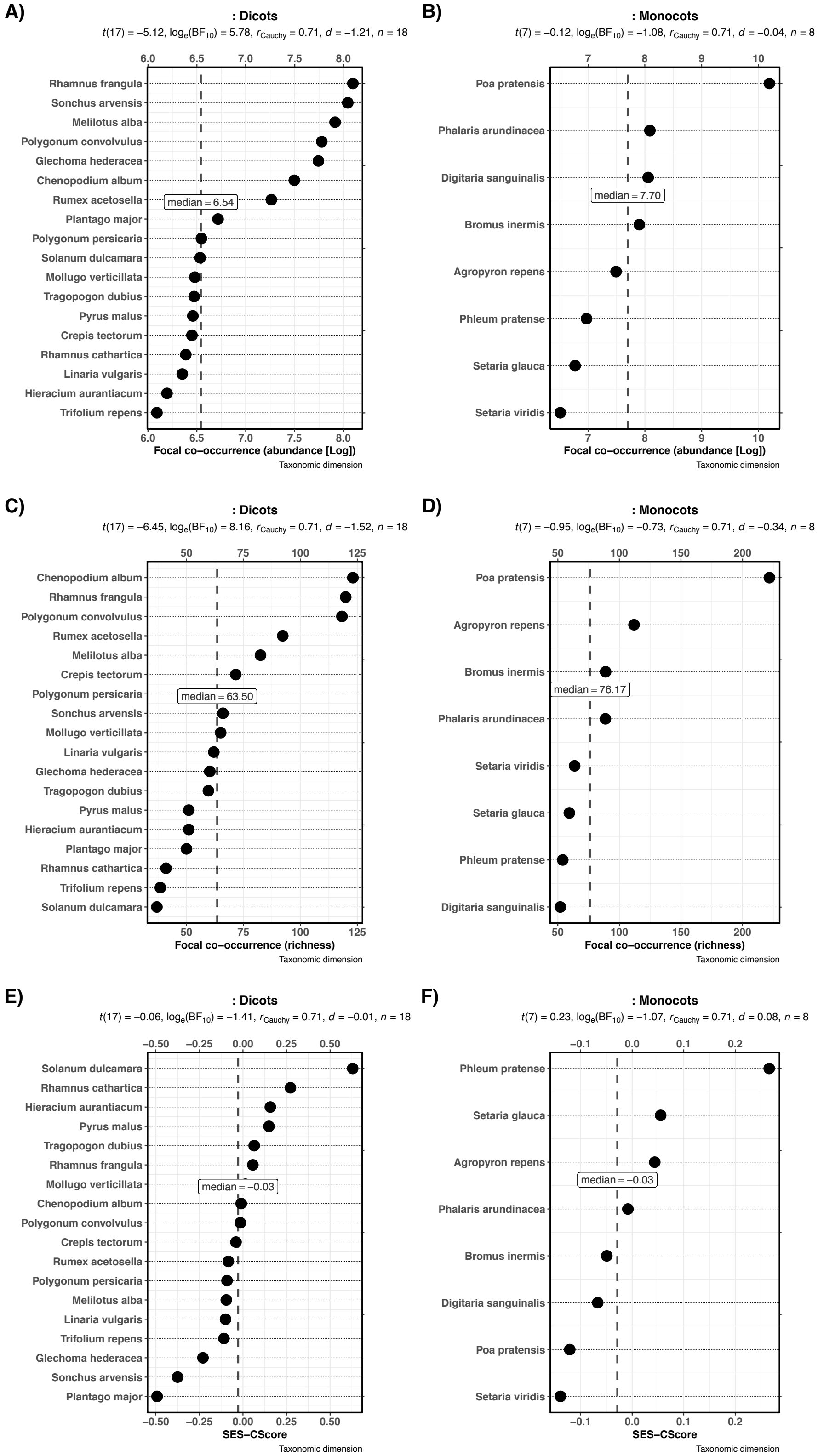
A)
: Dicots
t(17) = −5.12, log_e(BF_10) = 5.78, r_Cauchy = 0.71, d = −1.21, n = 18
Rhamnus frangula
Sonchus arvensis
Melilotus alba
Polygonum convolvulus
Glechoma hederacea
Chenopodium album
Rumex acetosella
Plantago major
Polygonum persicaria
Solanum dulcamara
Mollugo verticillata
Tragopogon dubius
Pyrus malus
Crepis tectorum
Rhamnus cathartica
Linaria vulgaris
Hieracium aurantiacum
Trifolium repens
median = 6.54
6.0 6.5 7.0 7.5 8.0
Focal co−occurrence (abundance [Log])
Taxonomic dimension
B)
: Monocots
t(7) = −0.12, log_e(BF_10) = −1.08, r_Cauchy = 0.71, d = −0.04, n = 8
Poa pratensis
Phalaris arundinacea
Digitaria sanguinalis
Bromus inermis
Agropyron repens
Phleum pratense
Setaria glauca
Setaria viridis
median = 7.70
7 8 9 10
Focal co−occurrence (abundance [Log])
Taxonomic dimension
C)
: Dicots
t(17) = −6.45, log_e(BF_10) = 8.16, r_Cauchy = 0.71, d = −1.52, n = 18
Chenopodium album
Rhamnus frangula
Polygonum convolvulus
Rumex acetosella
Melilotus alba
Crepis tectorum
Polygonum persicaria
Sonchus arvensis
Mollugo verticillata
Linaria vulgaris
Glechoma hederacea
Tragopogon dubius
Pyrus malus
Hieracium aurantiacum
Plantago major
Rhamnus cathartica
Trifolium repens
Solanum dulcamara
median = 63.50
50 75 100 125
Focal co−occurrence (richness)
Taxonomic dimension
D)
: Monocots
t(7) = −0.95, log_e(BF_10) = −0.73, r_Cauchy = 0.71, d = −0.34, n = 8
Poa pratensis
Agropyron repens
Bromus inermis
Phalaris arundinacea
Setaria viridis
Setaria glauca
Phleum pratense
Digitaria sanguinalis
median = 76.17
50 100 150 200
Focal co−occurrence (richness)
Taxonomic dimension
E)
: Dicots
t(17) = −0.06, log_e(BF_10) = −1.41, r_Cauchy = 0.71, d = −0.01, n = 18
Solanum dulcamara
Rhamnus cathartica
Hieracium aurantiacum
Pyrus malus
Tragopogon dubius
Rhamnus frangula
Mollugo verticillata
Chenopodium album
Polygonum convolvulus
Crepis tectorum
Rumex acetosella
Polygonum persicaria
Melilotus alba
Linaria vulgaris
Trifolium repens
Glechoma hederacea
Sonchus arvensis
Plantago major
median = −0.03
−0.50 −0.25 0.00 0.25 0.50
SES−CScore
Taxonomic dimension
F)
: Monocots
t(7) = 0.23, log_e(BF_10) = −1.07, r_Cauchy = 0.71, d = 0.08, n = 8
Phleum pratense
Setaria glauca
Agropyron repens
Phalaris arundinacea
Bromus inermis
Digitaria sanguinalis
Poa pratensis
Setaria viridis
median = −0.03
−0.1 0.0 0.1 0.2
SES−CScore
Taxonomic dimension

**Fig. S6**

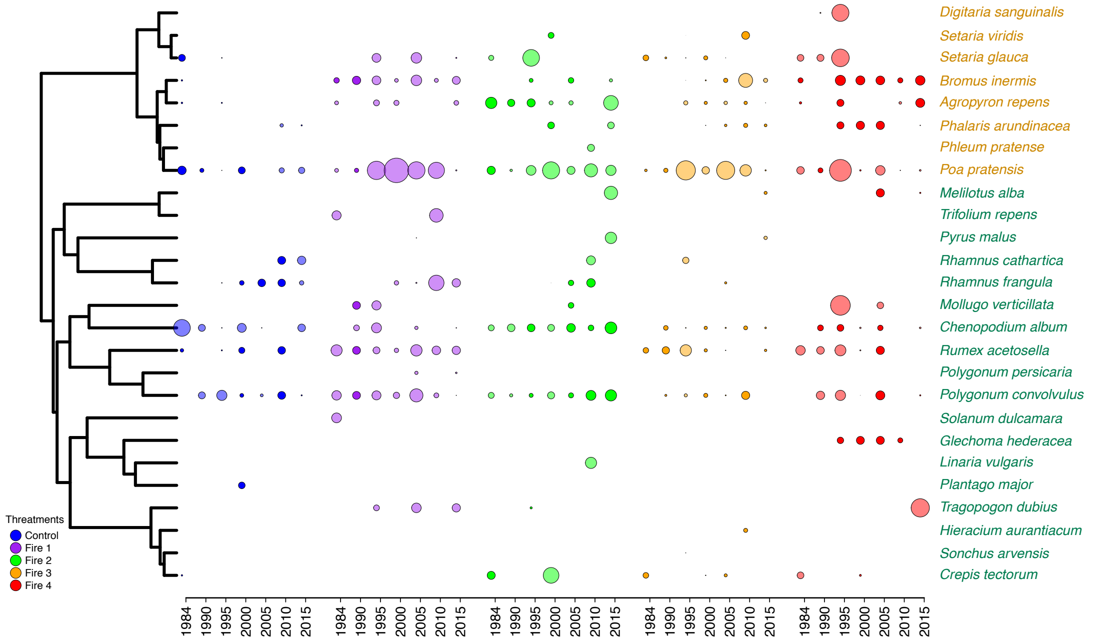
Digitaria sanguinalis
Setaria viridis
Setaria glauca
Bromus inermis
Agropyron repens
Phalaris arundinacea
Phleum pratense
Poa pratensis
Melilotus alba
Trifolium repens
Pyrus malus
Rhamnus cathartica
Rhamnus frangula
Mollugo verticillata
Chenopodium album
Rumex acetosella
Polygonum persicaria
Polygonum convolvulus
Solanum dulcamara
Glechoma hederacea
Linaria vulgaris
Plantago major
Tragopogon dubius
Hieracium aurantiacum
Sonchus arvensis
Crepis tectorum
Threatments
Control
Fire 1
Fire 2
Fire 3
Fire 4
1984 1990 1995 2000 2005 2010 2015 1984 1990 1995 2000 2005 2010 2015 1984 1990 1995 2000 2005 2010 2015 1984 1990 1995 2000 2005 2010 2015 1984 1990 1995 2000 2005 2010 2015

**Fig. S7**

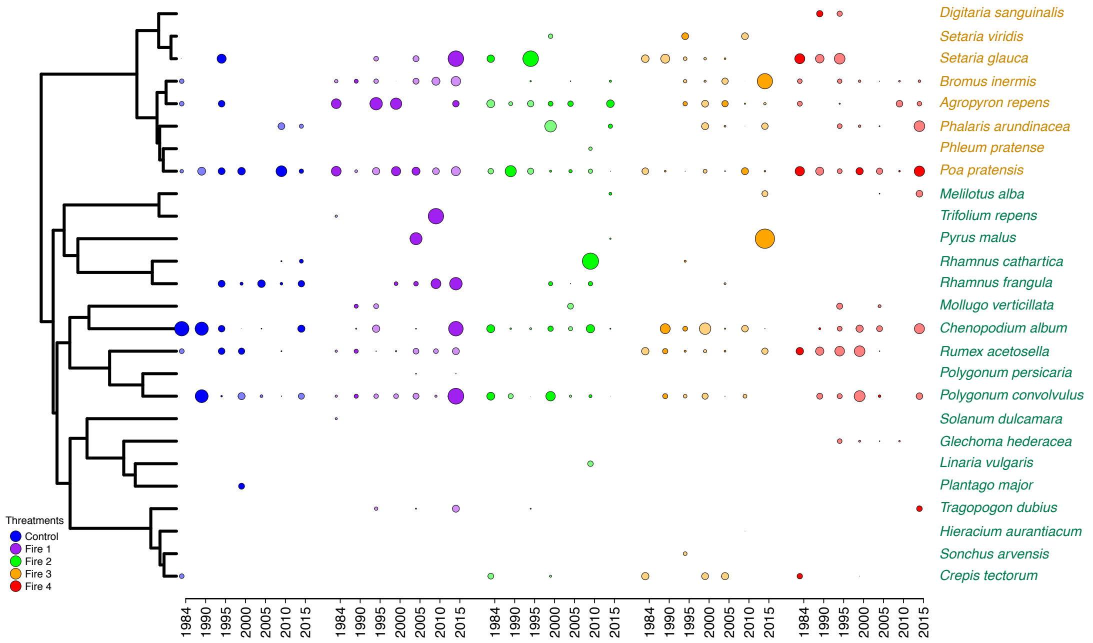
Digitaria sanguinalis
Setaria viridis
Setaria glauca
Bromus inermis
Agropyron repens
Phalaris arundinacea
Phleum pratense
Poa pratensis
Melilotus alba
Trifolium repens
Pyrus malus
Rhamnus cathartica
Rhamnus frangula
Mollugo verticillata
Chenopodium album
Rumex acetosella
Polygonum persicaria
Polygonum convolvulus
Solanum dulcamara
Glechoma hederacea
Linaria vulgaris
Plantago major
Tragopogon dubius
Hieracium aurantiacum
Sonchus arvensis
Crepis tectorum
Threatments
Control
Fire 1
Fire 2
Fire 3
Fire 4
1984 1990 1995 2000 2005 2010 2015 1984 1990 1995 2000 2005 2010 2015 1984 1990 1995 2000 2005 2010 2015 1984 1990 1995 2000 2005 2010 2015 1984 1990 1995 2000 2005 2010 2015
